# Structural evolution of the tissue-specific U2AF2 paralog and alternative splicing factor LS2

**DOI:** 10.1101/2020.08.15.252130

**Authors:** Ashish Ashok Kawale, J. Matthew Taliaferro, Hyun-Seo Kang, Christoph Hartmüller, Arie Geerlof, Ralf Stehle, Christopher Burge, Donald C. Rio, Michael Sattler

## Abstract

The *Drosophila melanogaster* LS2 protein is a tissue-specific paralog of U2AF2 that mediates testis-specific alternative splicing. In order to understand the structural mechanisms underlying the distinct RNA binding specificity we determined the solution structures of the LS2 RNA recognition motif (RRM) domains and characterized their interaction with *cis*-regulatory guanosine-rich RNA motifs found in intron regions upstream of alternatively spliced exons. We show that the guanosine-rich RNA adopts a G quadruplex (G4) fold *in vitro*. The LS2 tandem RRMs adopt canonical RRM folds that are connected by a 38-residue linker that harbors a small helical motif α_0_. The LS2 RRM2 domain and the α_0_ helix in the interdomain linker mediate interactions with the G4 RNA. The functional importance of these unique molecular features in LS2 is validated by mutational analysis *in vitro* and RNA splicing assays *in vivo*. RNA sequencing data confirm the enrichment of G4-forming LS2 target motifs near LS2-affected exons. Our data indicate a role of G quadruplex structures as *cis*-regulatory motifs in introns for the regulation of alternative splicing, that engage non-canonical interactions with a tandem RRM protein. These results highlight the intriguing molecular evolution of a tissue-specific splicing factor from its conserved U2AF2 paralog as a result of (retro-) gene duplication in *D. melanogaster*.

## Introduction

Alternative splicing is an essential process in post-transcriptional gene regulation in metazoans, leading to proteomic diversification without altering genome composition^1-4^. Alternative splicing is often dysregulated in human diseases including cancer^5,6^, neurodegenerative disorders, cardiovascular diseases, immunological and metabolic disorders^7^, reviewed in^8-10^. The regulation of alternative splicing is achieved via numerous interactions of splicing factors, i.e. RNA binding proteins (RBPs) with their cognate *cis*-regulatory elements ^11-15^. The net combinatorial effect of these interactions defines a “splicing code” that determines the pattern and the efficiency of the splicing reaction^16-20^. Although many alternative splicing factors and their interactions with *cis*-regulatory RNA elements have been identified, molecular and structural details of how similar or homologous RBPs acquire modulated specificities in order to orchestrate divergent functions are still poorly understood^12,21,22^.

U2 auxiliary factor (U2AF) is an essential splicing factor, which plays a key role in the assembly of the spliceosome by defining 3’ splice sites in eukaryotic introns^23^. U2AF is a highly conserved heterodimer found in all eukaryotes. The U2AF large subunit (U2AF2: hU2AF2 in *H. sapiens*, dU2AF2, in *D. melanogaster*) recognizes the intron poly-pyrimidine tract, whereas the small subunit (U2AF1: hU2AF1 in *H. sapiens*, dU2AF1 in *D. melanogaster*) interacts with the AG dinucleotide located at intron-exon junctions^23-31^. Structural and functional features of the U2AF heterodimer are well-studied^32-43^. The first two RNA recognition motifs (RRM1 and RRM2) in hU2AF2 recognize and interact with the poly-pyrimidine tract. In our previous studies, we have shown that the two domains exist in a conformational equilibrium between open and closed states, which plays a key role in adapting to diverse polypyrimidine tracts of different lengths and compositions^32,37,40,41,44^. Thus, U2AF can quantitatively link pyrimidine tract strength to the splicing initiation mediated by the conformation dynamics of the RNA binding domains.

LS2 is a *Drosophila* testis-specific paralog of dU2AF2, the large subunit of the U2AF heterodimer. The LS2 protein provides a fascinating paradigm for understanding evolutionary aspects of alternative splicing regulation. Even though LS2 shares high sequence homology to dU2AF2 (55% identity, 70% similarity at the protein sequence level), it exhibits very different RNA binding specificity (recognition of guanosine-vs. pyrimidine-rich RNAs), as well as splicing activity (alternative splicing repression vs. constitutive splicing activation)^45^ **(Fig. 1A, Supplementary Fig. 1)**. The expression of LS2 and LS2-regulated transcripts is found to be enriched in the testes and is linked to testes function, gamete production, and cellular regulation through phosphorylation. LS2 is proposed to perform its function by interacting with guanosine-rich sites flanking the poly-pyrimidine tract, but the underlying molecular mechanisms remains unclear^45^ **(Fig. 1B)**.

**Figure 1.**
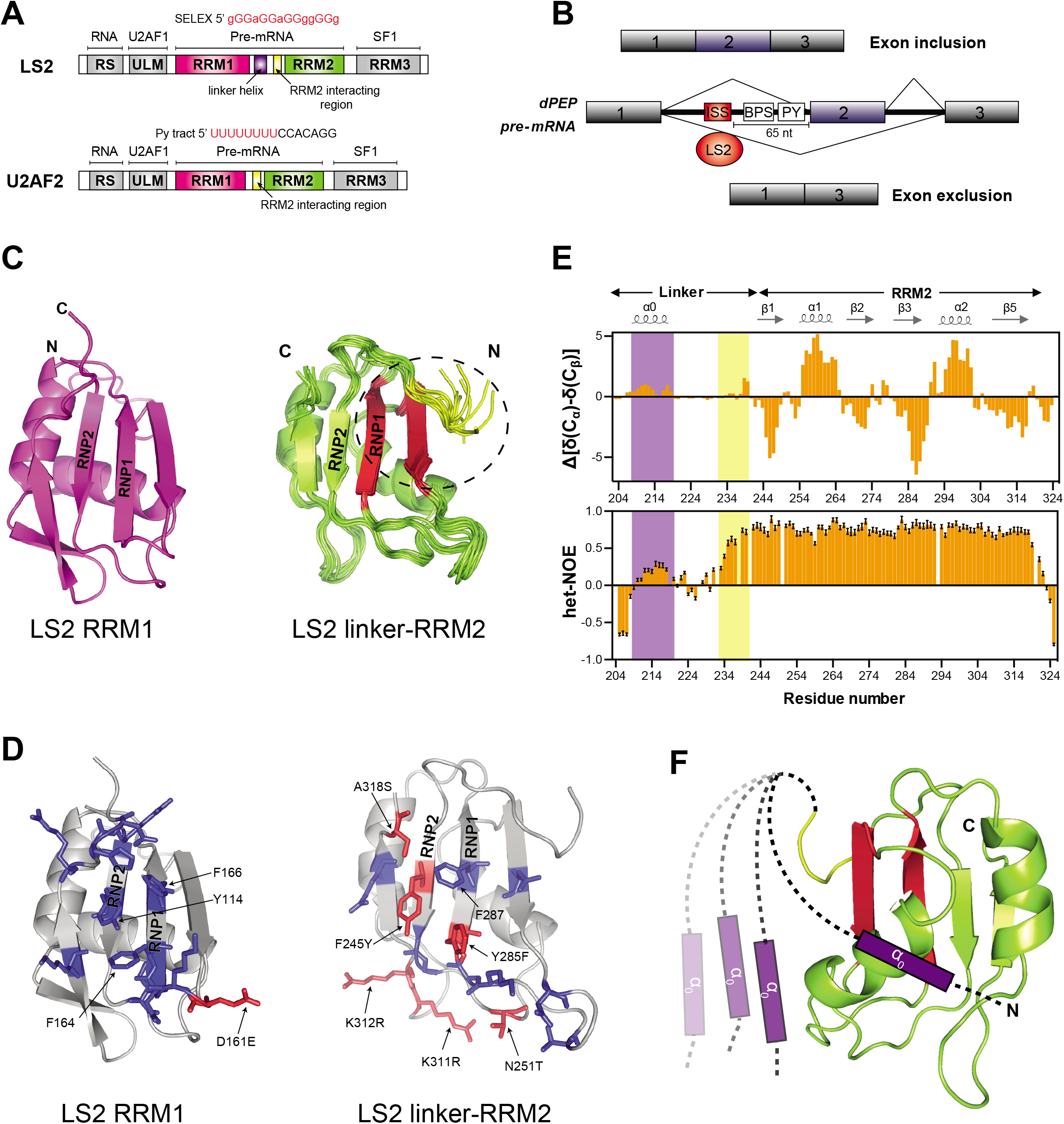
Overview of LS2 and U2AF2 domain architecture and NMR solution structures of LS2 RRM1 and linker-RRM2. **(A)** Schematic representations of LS2 and U2AF2 domain architecture and molecular functions. **(B)** The proposed role of LS2 in alternative splicing ^45^. **(C)** Lowest energy structure of RRM1 (backbone cartoon) (left) and the ensemble of the 10 lowest energy structures of the linker-RRM2 protein (right, only rigid residues 233–318 are shown). The interaction between RRM2 and C-terminal region of the linker is highlighted by a dotted eclipse and residues of RRM2 and C-terminal linker involved in the interaction are mapped in a red and yellow color, respectively. **(D)** Analysis of conservation of RNA binding residues highlighted in blue (conserved) and red (not conserved; annotated) on LS2RRM1 (left panel) and LS2 linker-RRM2 (right panel) structures, respectively in comparison to hU2AF2 RRM1,2 bound to poly-U RNA^36^ (see also **Supplementary Fig. 1**). The analysis indicates that the poly-U binding hU2AF2 residues are mostly conserved in LS2RRM1, whereas LS2RRM2 shows non-conservation of key poly-U binding residues including residues on RNP2 (Y245) and RNP1 (F285Y) sites, which mediates crucial RNA interactions. Conserved aromatic residues of the RNP sites are also annotated. **(E)** Chemical shift-derived secondary structure (top panel) and {^1^H}-^15^N heteronuclear NOE data for the LS2 linker-RRM2 in the free form. Secondary structure elements of the linker-RRM2 solution structure are shown on top. The more rigid α_0_ helix in the linker is highlighted with purple background, whereas the C-terminal semi-rigid region of the linker is marked with a yellow background. **(F)** Suggested interaction of the linker α_0_ helix with the RRM2 fold.

*Cis*-regulatory RNA sequence elements play important roles in the regulation of gene expression. These elements serve as binding sites for trans-acting RNA binding factors, though recent studies highlight their ability to function by adopting distinct secondary structure^22,46-49^. Guanosine-rich RNA motifs are found to be over-represented in the vicinity of splice sites of a growing number of genes^50^. Guanosine-rich RNA sequences can adopt higher order secondary structures called G-quadruplexes (G4) or G-quartets^50-54^. These structures are formed by stacked arrangements of G-quartets, in which four guanosines engage in Hoogsteen base-pairing, that *in vitro* are stabilized by alkali metal ions, such as potassium at high concentrations^55,56^. Recent studies highlight the importance of RNA G4 structures in important biochemical processes such as telomere maintenance, splicing, polyadenylation, RNA turnover, transcription, mRNA targeting and translation^54,57-72^. Moreover, G4 structures are linked to cancers and neurodegenerative diseases, such as Fragile X mental retardation syndrome and amyotrophic lateral sclerosis (ALS), by influencing actions of FMRP and hnRNP H splicing factors, respectively^54,73,74^. Notwithstanding the biological importance of G4, a comprehensive understanding of G4 structural features and their protein binding functional roles *in vivo* remain elusive^75^.

Given the high degree of primary sequence similarity, the distinct functional properties of LS2 and U2AF2 are surprising. It is important to understand whether functional differences between LS2 and U2AF2 stem from the structural differences of the LS2 and U2AF2 RNA binding domains. It is also unclear whether LS2 target RNAs can adopt a G4-fold and whether these structures play any role in the LS2-mediated splicing regulation. In order to answer these questions, we have solved the solution NMR structures of LS2 RNA binding domains and characterized the guanosine-rich LS2 RNA ligands. Structural characterization reveals that, despite retaining a canonical RRM structure, the LS2 RRMs have evolved to recognize the G4-RNA fold. RRM2 provides the specificity for this interaction, while additional binding contributions are provided by RRM1 and a unique short helical motif in the RRM1,2 linker. The LS2-G4 interaction presented in this study shows that the RRM fold and flanking regions can recognize G4 structures, distinct to the common recognition of single-stranded nucleic acid sequences. Our results add another dimension to the versatile RNA binding nature of the RRM domain.

## Results

### Structural characterization of LS2 RNA-binding domains

Given the distinct RNA binding sequence preferences of LS2 vs U2AF2, we first focused on the minimal RNA binding region comprising the tandem RRM1 and RRM2 domains, which were previously shown to mediate RNA binding by LS2^45^ **(Fig.1A, Supplementary Fig. 1)**. As the tandem RRM1,2 construct was not sufficiently stable for structural studies, we employed a divide-and-conquer approach by analyzing the individual RRM1, RRM2 domains, as well as a region comprising RRM2 and the preceding linker. The NMR structures of RRM1 and RRM2 show that both LS2 RRMs exhibit a canonical βαββαβ fold, characterized by four antiparallel β-strands packed against two α-helices (**Fig. 1C)**. NMR ^15^N relaxation data also corroborate the canonical folds of LS2 RRM domains (**Supplementary Fig. 2, Supplementary Information**). We first compared residues involved in poly-pyrimidine-tract recognition in hU2AF2 with those in dU2AF2 and LS2. This analysis shows that these residues are strictly conserved in dU2AF2 and mostly conserved in LS2 RRM1, whereas some minor variations are seen in LS2 RRM2 (**Fig. 1D, Supplementary Fig. 1**). This observation is consistent with analysis of hU2AF2 where specificity for the uridine nucleotides is provided by the RRM2, whereas RRM1 is more promiscuous in nature^34^. Potentially, the change of two lysine side chains by arginine could affect the specific RNA recognition by RRM2.

To map potential domain interactions within LS2 RRM1,2, we compared the ^1^H-^15^N NMR correlation spectra of various protein fragments of LS2 (**Supplementary Fig. 3A**). NMR chemical shifts of residues in the RRM1 domain alone are very similar compared to those in the tandem domain constructs (RRM1,2 or RRM1,2 Δlinker). In contrast, for RRM2, significant chemical shift differences are observed, especially for residues located in strands β2 and β3 (RNP1) comparing RRM2 vs RRM1,2 (**Supplementary Fig. 3A, B)**. These chemical shift differences are not observed when the spectrum of the linker-RRM2 construct is compared to RRM1,2, while they are seen when comparing RRM1,2 and the RRM1,2 Δ linker protein. This indicates that the linker residues interact with the β-sheet of RRM2, in proximity to the RNP sites. The NMR structure of the linker-RRM2 protein fragment indeed reveals this interaction where the hydrophobic residues in the C-terminal region (residues 233–237) of the linker interact with the β2 and β3 strands of RRM2 **(Fig. 1C**. The multiple sequence alignment also reveals the higher degree of conservation of residue 233–237 in the C-terminal region of the RRM1-RRM2 interdomain linker throughout the U2AF2 paralogs **(Supplementary Fig. 1)**. This interaction has been recently also reported to modulate RNA binding by hU2AF2^44^.

Interestingly, the RRM1,2 linker in LS2 is significantly longer than in the human or Drosophila U2AF2 orthologues with an N-terminal extension of 28 amino acids (residues 205–232) **(Supplementary Fig. 1**). NMR relaxation data show that the N-terminal extension, and also C-terminal region of the RRM1,2 linker preceding the RRM2 mentioned above, exhibit reduced flexibility indicating some structural propensity **(Fig. 1E; Supplementary Fig. 4A)**. NMR secondary chemical shifts indicate that residue 210–218 in the LS2-specific N-terminal region sample an α-helical conformation **(Fig. 1E)**. Further analysis revealed that this helical region does not (transiently) interact with the RRM1 domain as chemical shifts of the helical residues in the linker-RRM2 protein fragment match very well to the RRM1,2 protein, but are distinct in the RRM1-linker fragment that lacks RRM2 (**Supplementary Fig. 3A, 3B; Supplementary Fig. 4B)**. This suggests that the N-terminal helical region provides an additional dynamic contact with RRM2, in addition to the C-terminal linker region. NMR analysis of a shortened linker-RRM2 protein fragment lacking this helix indeed shows chemical shift changes mainly for helix α1 and a loop preceding the β5 strand of RRM2 (**Supplementary Fig. 4C**). This indicates that the LS2-specific linker helix could modulate or contribute to RNA binding of RRM2. Small angle X-ray scattering (SAXS) data for RRM1,2 indicate the presence of an ensemble of conformations, including compact and non-compact states (**Supplementary Fig. 4D**) very similar to what has been observed previously for hU2AF2^32,37^. This indicates that the two RRM domains in LS2 are connected by a flexible linker and can sample a range of relative arrangements in solution that could potentially play a role for its function.

In summary, the LS2 RRM1,2 region comprises two canonical RRM domains that are connected by a semiflexible linker harboring two stretches with reduced flexibility. One region directly preceding and binding to the surface of RRM2 (**Fig. 1C**) resembles hU2AF2, while the second one following RRM1 form a small helical motif that transiently interacts with RRM2 (**Fig. 1E, F**). Both regions are highly conserved throughout the LS2 homologs from various Drosophila species (**Supplementary Fig. 4E**).

### LS2 binds to a G-quadruplex-forming guanosine-rich RNA motif

LS2 has been previously shown to bind to G-rich RNAs^45^. Given that guanosine-rich oligonucleotides can adopt G quadruplex (G4) structures, we studied the conformation of G-rich RNA ligands of LS2. We used the 21-mer (5’-GGGGAGGAGGGGGGCGUAUGA-3’) RNA, which was designed according to SELEX data and has been shown to bind to purified LS2 protein^45^. Notably, a circular dichroism spectrum of this RNA shows a positive absorption at 265nm and negative absorption at 240nm. This indicates the formation of a parallel-stranded G4 structure in the presence of KCl or NaCl, as well as in the absence of salt (**Supplementary Fig. 5A**)^76-78^. The ^1^H 1D NMR spectrum shows numerous imino signals at 10–12 ppm indicative of Hoogsteen base-pairing, which is consistent with the formation of a G4 structure in the presence of 150 mM KCl (**Supplementary Fig. 5B**)^56,79-81^. As is often observed with G4 structures, the NMR spectrum shows the presence of multiple heterogeneous and dynamic conformations, indicated by multiple imino resonances with severe line-broadening centered around 11 ppm^57,79,81^. By screening a number of monovalent ion salt conditions, we found that this RNA adopts a homogeneous G4 conformation in the presence of low K^+^ concentrations, characterized by sharp imino resonances around 10–12 ppm in a ^1^H 1D NMR spectrum (**Supplementary Fig. 5C**).

To further characterize the RNA G4 conformation at low K^+^ concentration, we used ^1^H NMR spectroscopy. At 278 K, a ^1^H 1D NMR spectrum shows the presence of at least eight sharp and well resolved resonances in the imino region (10–12 ppm) **(Fig 2A)**. These resonances correspond to the guanosine imino protons as confirmed by characteristic ^15^N chemical shifts observed in a natural abundance ^1^H-^15^N HSQC spectrum (**Supplementary Fig. 5D**). The variations in signal intensities and linewidths indicate the presence of additional overlapping resonances, which become more resolved with reduced linewidths observed at increasing temperature. At 318 K, eight resonances with comparable linewidths (annotated as 1, 2, 3, 4, 5 and 8 in spectra) and two others (annotated as 6, 7) with two-fold higher intensity are observed (**Fig. 2A**). This is consistent with the presence of 12 imino protons, as is expected for a three-planar G4 structure. Two broader signals upfield of the imino region (indicated by “*”) correspond to the purine amino group (**Supplementary Fig. 5D**). The fact that these amino proton resonances (which are typically highly exchangeable with solvent) are observed is indicative of a slow exchange of the amino protons and their involvement in hydrogen bonding, consistent with the formation of a A:(G:G:G:G):A hexad structure **(Fig. 2B)**^81-83^. The positive absorption around 305 nm in the CD spectrum of RNA in the presence of low K^+^ concentrations is also indicative of hexad formation^82^ (**Supplementary Fig. 5E**). Hexad formation has been linked to GGA motif-containing nucleic acid sequences^81,82,84^. CD thermal melting studies further corroborated the formation of well-folded G-quadruplex species with the melting temperature around 53 °C (**Supplementary Fig. 5F**).

**Figure 2.**
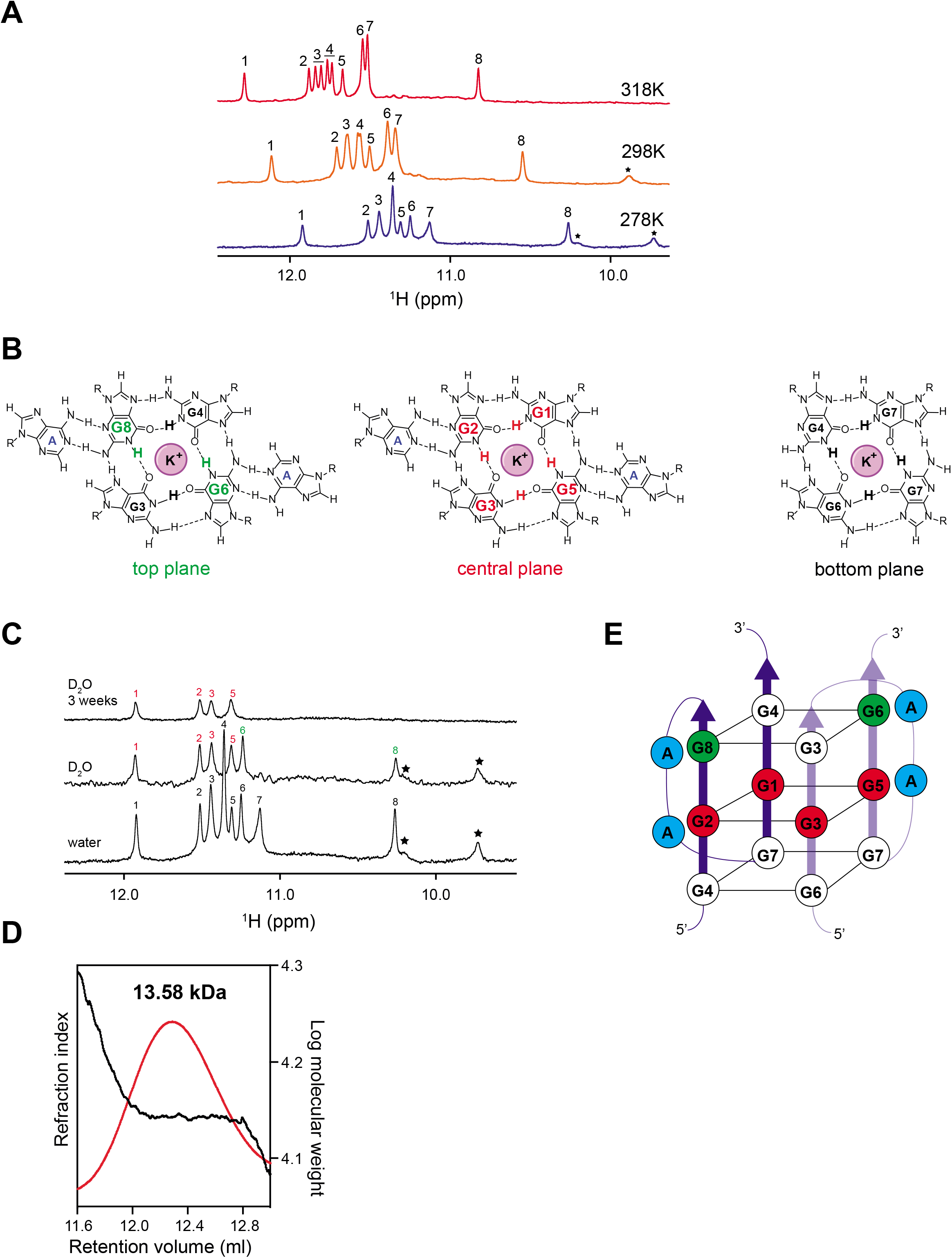
LS2 target RNA adopts G4 fold with three planar, parallel and dimeric topology. **(A)** ^1^H 1D NMR spectra of the imino region of the 21-mer RNA in the presence of 5 mM KCl at different temperatures recorded on 500 μM RNA sample at 600 MHz. The imino resonances are numbered 1–8 depending upon their position in the ^1^H NMR spectrum at 278K. Purine amino resonances are indicated by “*”. **(B)** Chemical structure of the proposed G-quadruplex topology with A:(G:G:G:G):A hexads (top and central plane) and a G:G:G:G tetrad (bottom plane). **(C)** H/D exchange experiments of the 21-mer RNA in the presence of 5 mm KCl at various time intervals recorded on a 500 μM RNA sample at 600 MHz, 278 K. The spectrum after three weeks of D_2_O exchange was recorded with 128 times higher number of scans and is scaled for a better comparison. **(D)** Static light scattering for the homogeneous conformation of the 21-mer RNA. The main peak of interest is shown with a plot of refractive index vs. log molecular weight plot. The calculated molecular mass is 13.58 kDa which is approximately double of the theoretical molecular of 7.23 kDa for RNA indicating dimeric RNA population in the solution. **(E)** Schematic representation of the dimeric RNA G4 formed by 21-mer G4 in the presence of 5mM KCl. Top two planes form A:(G:G:G:G):A hexads, whereas the bottom plane forms G:G:G:G tetrads. Guanine nucleotides from the top plane, which are involved in the hexad formation are colored in the green whereas guanines from the central plane are colored in red. Adenosines involved in the hexads formation are colored in blue. Two molecules of RNA are indicated in a blue and a light blue color, respectively.

In order to confirm the presence of a three planar G4 structure and hexad formation, we performed H/D exchange studies monitored by ^1^H NMR (**Fig. 2C**). This reveals that six imino resonances are protected against solvent exchange. Out of these imino signals four remain detectable even after three weeks in D2O buffer. In addition, two amino resonances also show partial protection to solvent exchange. This further supports the presence of a three-planar G4 structure, where the central plane, which is sandwiched between two other G-quartets, is expected to be more protected against solvent exchange. Formation of the hexad structure is ascertained by the partial solvent protection of two imino protons possibly due to hydrogen bonding with neighboring adenosine nucleotides **(Fig. 2B**).

To determine the molecular weight and stoichiometry of the G4 RNA we performed size exclusion chromatography coupled with static light scattering. Static Light scattering data corroborates the monodispersity of the G4 RNA structure with a molecular mass of 13.58 kDa, which is approximately twice the theoretical molecular weight of RNA (7.23 kDa) (**Fig. 2D**), and thus indicates a dimeric G4-fold. The imino NOEs obtained from the ^1^H-^1^H imino NOESY spectra are in line with our proposed model of the three planar, dimeric RNA G4 structure (**Supplementary Fig. 6, Supplementary Information**)^56,80^. In summary, our data indicate that the 21-mer G-rich RNA adopts a dimeric 3-planar G quadruplex with a hexad made up from GGA motifs. **(Fig. 2E**).

### LS2 specifically recognizes the G4 RNA fold

We next performed protein-RNA titration NMR experiments to monitor the complex formation and map the binding interface. Changes in backbone amide chemical shifts of the LS2 RRM1,2 tandem domains upon addition of RNA were monitored by ^1^H-^15^N HSQC experiments, whereas G4 formation was assessed by monitoring imino signals in ^1^H 1D NMR experiments. Upon addition of RNA the LS2 RRM1,2 protein starts to aggregate forming gel-like aggregates, leading to severe line-broadening for the amide signals (**Supplementary Fig. 7A**). A similar behavior is observed for RNA titrations with LS2 RRM1. In contrast, RNA titration with the LS2 RRM2 or linker-RRM2 constructs do not show any visible aggregates. Upon addition of RNA to individual RRM1 and RRM2 domains line broadening and some chemical shift changes are observed, indicating protein-RNA interaction in the intermediate exchange regime and/or in multiple registers (**Supplementary Fig. 7B, D**).

To map the RNA interfaces, we therefore analyzed the effects of the addition of the 21-mer RNA to the individual RRM1 and RRM2 domains. Sub-stoichiometric addition of the RNA to RRM1 induces linebroadening for numerous residues. Significant chemical shift perturbations (CSPs) are seen, especially for residues in the β5 strand, which predominantly harbors positively charged residues (K192, R194, R195). CSPs are also observed for the C-terminal tail, containing aromatic residues (H197 and Y199), as well as the consensus RNP2 motif (**Supplementary Fig. 7C**). No significant spectral changes are observed for residues of the RNP1 motif. In contrast, addition of RNA to RRM2 results in chemical shift perturbations for residues located in both RNP motifs and the neighboring β-strands (**Supplementary Fig. 7E**). These observations indicate that RRM2 interacts with RNA using the canonical RNP sites. The linker-RRM2 protein fragment shows similar spectral changes for RRM2 (**Fig. 3A–C**). However, additional CSPs are observed for residues in the LS2-specific α_0_ helix in the RRM1-RRM2 linker This argues that these residues are affected by the presence of RNA by direct contacts, as residues both in RRM2 and the α_0_ helix exhibit line broadening. Isothermal titration calorimetry data show that the dissociation constant (*K_d_*) of the linker-RRM2-RNA interaction is in the low micromolar range (**Fig. 3D**). This is in line with the NMR titration data showing line broadening upon RNA binding. This affinity is consistent with the *K_d_* value of 1.9 μM observed for full-length LS2-poly G RNA binding^45^. These data indicate that the main RNA binding regions involve RRM2 and the α_0_ helix in the RRM1-RRM2 linker. RRM1 may mediate additional non-specific contributions to the RNA interaction, as indicated in the NMR titrations of RRM1,2 (**Supplementary Fig.7A**).

**Figure 3.**
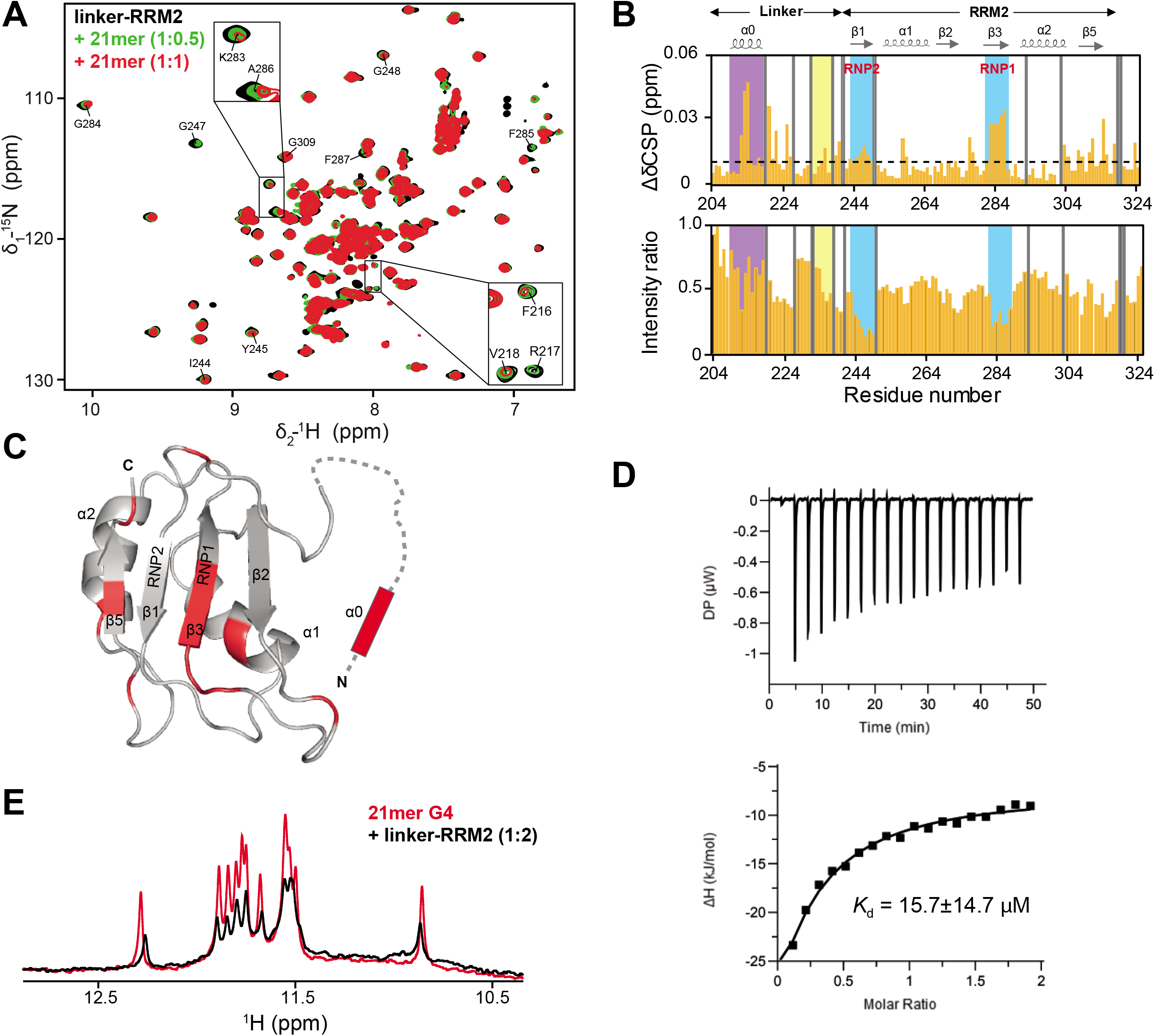
LS2 linker-RRM2 interacts with the G4 RNA structure. **(A)** The overlay of the ^1^H-^15^N HSQC spectra of the LS2 linker-RRM2 construct, in the absence (black) and in the presence of 0.5 molar excess (green) and one molar excess (red) of the 21-mer G4 RNA. Selected residues from the RNP sites and the linker helix α_0_ are annotated. **(B)** Backbone amide chemical-shift perturbations (CSP, Δδ) (top panel) and intensity change (bottom panel) for the linker-RRM2 residues upon 21-mer RNA binding. The secondary structure of the linker-RRM2 construct is shown at the top. Grey bars in the plot indicate missing resonance assignments or proline residues. The RNP1/2 sites are highlighted with cyan background, the linker-helix is marked with purple background, residues at the C-terminal region of the linker that interact with RRM2 are highlighted by yellow background. **(C)** Residues with CSPs greater than twice the standard deviation are mapped in red onto the solution NMR structure of the linker-RRM2 along with the α_0_ linker-helix. **(D)** ITC of LS2 linker-RRM2 binding to 21-mer RNA. The top panel shows the raw calorimetric whereas bottom panel indicates binding isotherms describing the complex formation. **(E)** A ^1^H 1D NMR spectra of 21-mer RNA in 7 mM potassium phosphate, pH 6.5, 5 mM DTT in the absence (black) and in the presence of the two-molar excess of the linker-RRM2 (red), showing linker-RRM2 and G4 interaction.

We hypothesized that the observed relatively small chemical shift perturbations in comparison to typical protein-single-stranded RNA complex formation may indicate the presence of folded RNA structure in the complex. To address whether LS2 recognizes the G4 RNA structure formed by RNA, we performed reverse titration experiments, in which the linker-RRM2 was titrated to G4 RNA in buffer conditions, which favor a homogeneous G4 conformation (low K^+^ concentration) (**Fig. 3E**). Upon successive addition of the protein the G4 specific imino resonances show line broadening and minor chemical shift perturbations as monitored by ^1^H NMR. Line broadening of imino resonances in the presence of protein might also imply that protein may bind to smaller proportion of single-stranded RNA and thus indirectly favor G4 unfolding by shifting the equilibrium towards single-stranded RNA form as observed for hnRNP F ^85^. However, the 1:1 molar binding ratio that we have observed by ITC rule out this possibility.

These data strongly argue that protein recognizes the G4 RNA conformation. Interestingly, upon titration of linker-RRM2 protein to RNA in buffer conditions, in which the RNA adopts a heterogeneous mixture of conformations, no significant spectral changes are observed, indicating that no or only very little protein-RNA complex is formed (**Fig. 4A**). It appears that a heterogeneous G4 population is kinetically trapped and cannot be simply reverted to a homogenous population by buffer exchange. However, upon unfolding and subsequent slow cooling (from 95 °C to room temperature) in a low K^+^ concentration buffer the RNA refolded to a more homogeneous RNA species (**Fig. 4B**). This allowed us to directly observe the linker-RRM2-RNA complex formation by unfolding and slow cooling (**Fig. 4C**). These data suggest that the linker-RRM2 protein specifically recognizes the three planar G4 folded hexad RNA conformation that predominates at low K^+^ concentrations.

**Figure 4.**
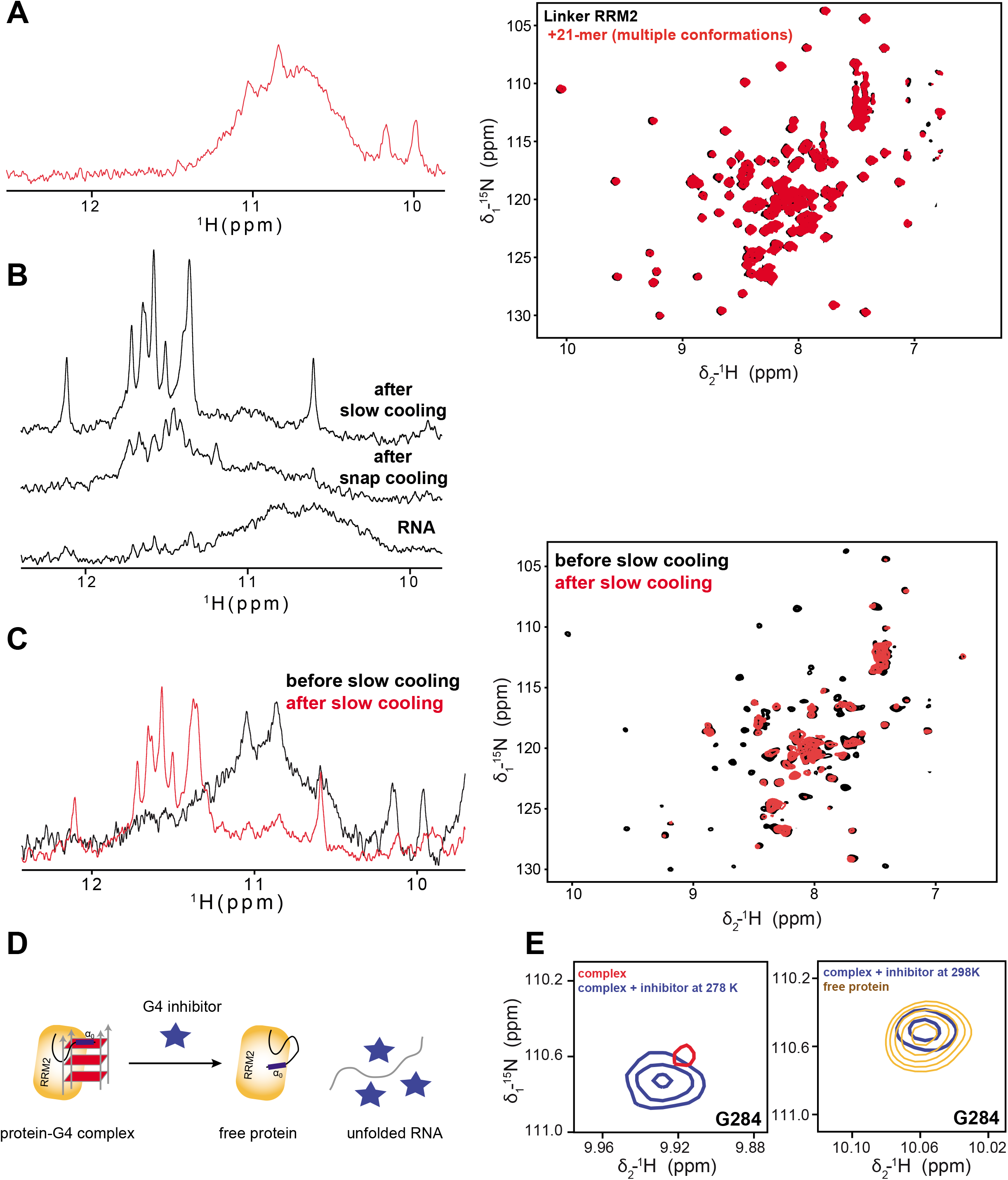
LS2 linker-RRM2 interacts specifically with homogeneous conformation of poly G RNA. **(A)** ^1^H 1D NMR spectrum of the imino region of the 21-mer RNA used for NMR titration showing the heterogeneous RNA G4 population in 20 mM MES buffer 6.5, 50 mM KCl, 5 mM DTT at 298 K and 500 MHz (top panel). The overlay of ^1^H-^15^N spectra of RRM2 in the absence of RNA (black) and in the presence of two molar excess (red) of 21-mer RNA heterogeneous G4 population (bottom panel). **(B)** ^1^H NMR spectra of the imino region of RNA with heterogeneous G4 population after dialysis in 7 mM potassium phosphate buffer pH 6.5 and after unfolding and refolding via the snap cooling (heating at 95 °C followed by incubation at 4°C) and the slow cooling (heating at 95 °C followed by incubation at RT). **(C)** ^1^H 1D NMR spectra of the imino region showing the transition of the 21-mer RNA from heterogeneous to homogeneous population after slow cooling (top panel) in the protein-RNA complex. The overlay of ^1^H-^15^N spectra of linker-RRM2 in the presence of 21-mer heterogeneous population i.e. before slow cooling (red) and in the presence of homogeneous population i.e. after slow cooling (green). **(D)** Schematic representation of experimental design to check the linker-RRM2-21-mer complex disruption in the presence of the G4-specific inhibitor (TMPyP4). **(E)** A representative NMR signal indicating the disruption of the linker-RRM2-21-mer complex disruption upon addition of G4 specific inhibitor (TMPyP4). The left panel shows the increased signal intensity and chemical shift changes indicating the free protein for Gly284 in the presence of the inhibitor. The right panel shows that the chemical shift of the Gly284 amide after overnight incubation with the inhibitor matches with that of the free protein. Note, that the spectra shown at the left and right were recorded at 278 and 298 K, respectively, for technical reasons (see **Supplementary Fig. 8**).

The line broadening of the G4-specific imino resonances associated with protein binding renders it challenging to assess by NMR whether the G4-fold is fully maintained in the complex. In order to further confirm this, we used an RNA G4 structure specific binder TMPyP4, a cationic porphyrin ring, that has been previously reported to destabilize RNA G quadruplexes ^62,86,87^. A ^1^H 1D NMR spectrum recorded for the LS2-RNA complex in the presence of TMPyP4 shows the disappearance of the G4 specific imino resonances consistent with the inhibition of G4 RNA formation (**Supplementary Fig. 8A**). The ^1^H-^15^N HSQC spectrum of the LS2 linker-RRM2-G4 RNA complex exhibits line broadening of most of the signals, as mentioned above. Notably, in the presence of the G4 inhibitor the NMR signals of the unbound linker-RRM2 are observed, indicating that inhibition of the G4 RNA conformation by the inhibitor leads to an inhibition of the protein-RNA interaction (**Fig. 4D, E; Supplementary Fig. 8B**). Further incubation of the complex at room temperature overnight resulted in complete disruption of the complex, as seen by the sharp linker-RRM2 resonances indicative of the free protein **(Supplementary Fig. 8C**). These data indicate that the LS2 linker-RRM2 protein specifically recognizes the G4 structure and does not bind to single-stranded form of guanosine-rich RNA. Similarly, we also observed specificity of isolated LS2 RRM2 towards 21-mer specific G4 structure by testing binding in the presence of various shorter oligonucleotides **(Supplementary Fig. 9,10; Supplementary information**).

To confirm and validate the LS2 regions involved in RNA we performed mutational analysis of residues in LS2 that are implicated in RNA binding and monitored the protein-RNA interactions by NMR. To abolish RNA binding contributions from RRM2, RRM1,2^F285D, F287D^ was generated where two conserved residues from RNP1 sequence motif are altered. This double mutant abolished RNA binding by RRM2, while maintaining the RNA binding contribution from RRM1 and the linker (**Supplementary Fig. 11A**). Similarly, to eliminate RNA binding from the α_0_ helix in the RRM1-RRM2 linker (H197–P233) we replaced this region by a glycine-glycine-serine-rich (GGS) polypeptide of identical length to introduce a random coil conformation for these residues. The ^1^H-^15^N HSQC NMR spectrum of the GS_linker mutant shows that both RRM domains remain fully folded and that RNA interactions by the RRM domains is not affected. In contrast, the RNA binding contribution from the linker region is no longer present (**Supplementary Fig. 11B**). We also tested a RRM1,2^K192E, R194E^ double mutant that affects the RRM1 β5 strand. This double-mutant leads to unfolding of the RRM1 domain and thereby, prevents RNA binding of RRM1 without altering the RNA binding contributions from RRM2 and the linker (**Supplementary Fig. 11C**).

### Functional analysis of the splicing regulatory activity of LS2 mutants

To test the role LS2 RNA binding to regulate alternative splicing *in vivo*, we created *Drosophila* S2 cell lines that stably and inducibly express wild type or mutant versions of LS2 **(Fig. 5A)**. S2 cells do not express endogenous LS2. Therefore, these wildtype and mutant transgene versions of LS2 were the sole contributors of cellular LS2 protein. The mutant versions of LS2 were based on amino acid residues that we identified as critical for RNA binding, including mutants we named RRM1_beta5 (K192E, R194E), RRM2_RNP1 (F285D, F287D), and GS_linker (H197–P233 replaced with glycine-glycine-serine motifs). In addition, we included two mutants that are defective in forming protein-protein interactions with other splicing factors. These include U2AF1_int (W62A, D63A, I64A) and SF1_int (E370K, D371K) mutants that lack the ability to interact with dU2AF1 and SF1, respectively (**Supplementary Fig. 12A**)^39,45^. LS2 is known to strongly interact with dU2AF1, and this interaction is required for high affinity binding of LS2 to RNA^45^. However, the consequence of the LS2 interaction with SF1 is unknown. To facilitate quantification of each mutant using immunoblotting and to allow determination of the subcellular localization of LS2, each mutant was expressed as a GFP fusion protein.

**Figure 5.**
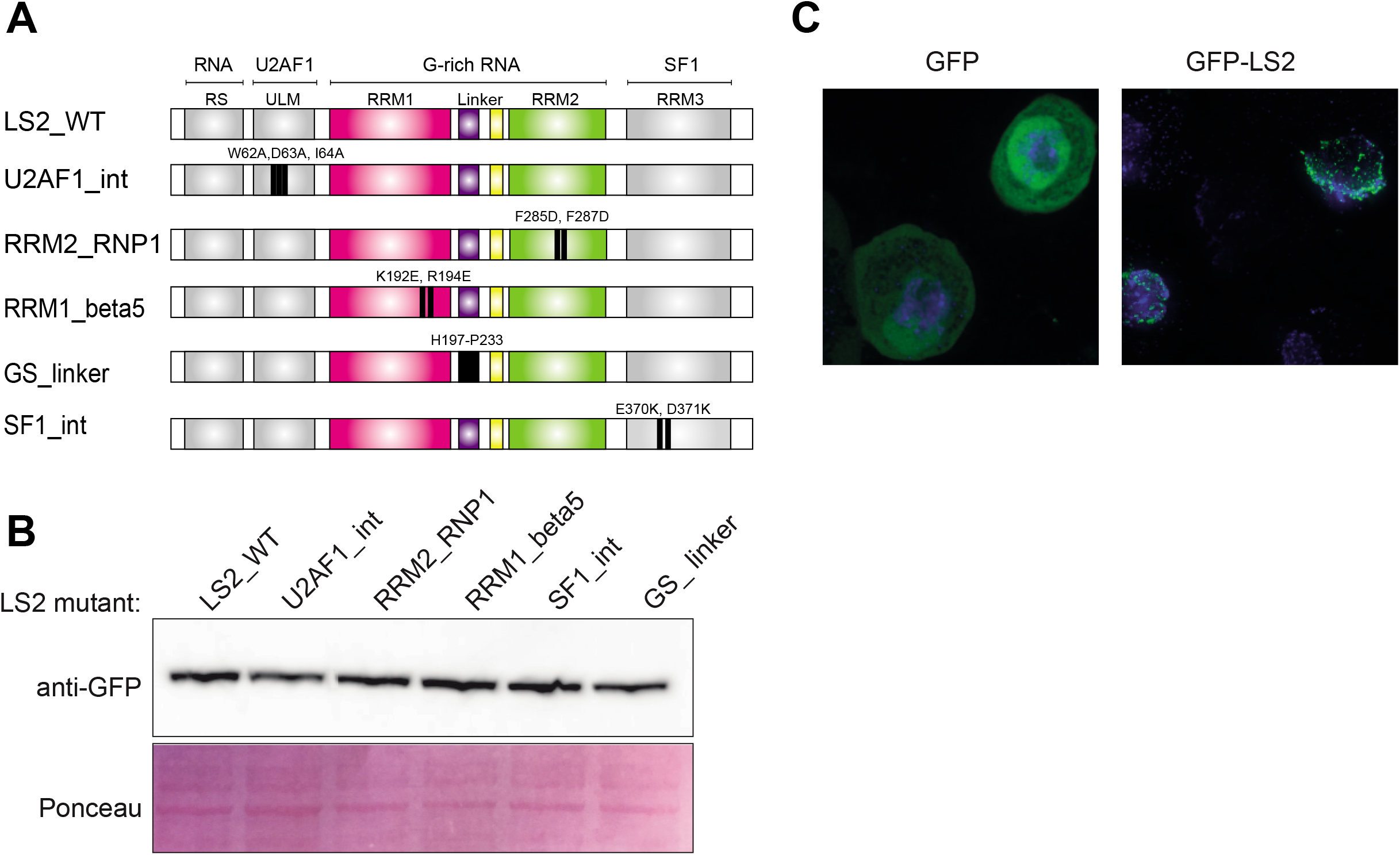
Expression of LS2 mutants in *Drosophila* S2 cells. **(A)** Diagram of locations of mutations (colored in black) in expressed LS2 mutants (Also, see **Supplementary Fig. 9**). **(B)** S2 cell lines with stably integrated, copper-inducible GFP-LS2 constructs were made. Upon induction with copper, the expression level of each mutant was assayed by immunoblotting with an anti-GFP antibody. **(C)** Fluorescence microscopy of GFP-LS2 fusions. While GFP alone (left) is localized throughout the cell, GFP-LS2 fusions are restricted to punctate bodies within the nucleus.

The expression of wild type and mutant LS2 proteins was induced for 53 hrs. Control cells expressing GFP alone were also induced as a negative control. To confirm that the mutants were expressed at comparable levels, lysates from each cell line were analyzed by immunoblotting using an anti-GFP antibody **(Fig. 5B)**. The subcellular localization of LS2 was examined using fluorescence microscopy. Interestingly, although GFP alone was distributed relatively evenly throughout the cell, LS2-GFP fusions were restricted to the nucleus and displayed defined, punctate foci **(Fig. 5C**), consistent with the known localization patterns of other splicing factors ^88^. RNA from each cell line was then collected in triplicate and profiled using high-throughput sequencing of Illumina RNA-seq libraries.

Quantification of LS2 RNA from each line again showed that wild type LS2 and the mutants were approximately equally expressed **(Supplementary Fig. 12B**). We quantified the levels of inclusion for alternatively spliced exons in each sample as percent spliced in (PSI) values. PSI values were highly correlated between replicates from the mutant RNA samples, but less so across mutants, indicating the high reproducibility of the data **(Supplementary Fig. 12C**). Endogenous LS2 protein is very poorly expressed in S2 cells **(Supplementary Fig. 12B**). We therefore identified alternative exons that were regulated by LS2 in S2 cells by comparing (PSI) values for exons in the GFP and wild type LS2-GFP lines. This analysis revealed 293 alternative exons that were significantly more excluded in the wild type LS2-GFP cells compared to GFP control cells and 164 exons that were more included, consistent with LS2 generally acting as a splicing repressor **(Fig. 6A**) ^45^.

**Figure 6.**
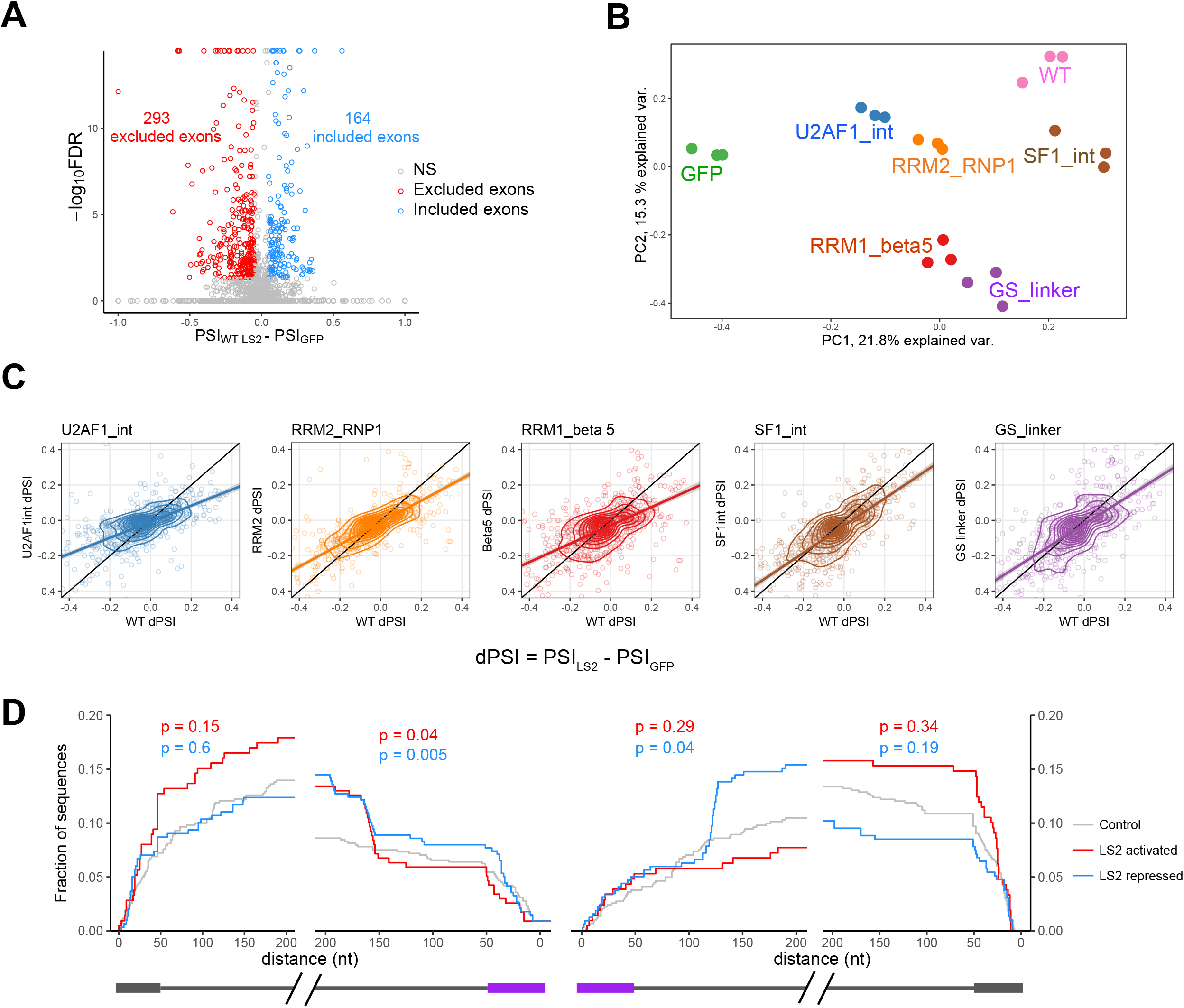
Alternative splicing regulatory activity of LS2 mutants. **(A)** The level of inclusion (percent spliced in or PSI) of alternative exons was calculated and compared in cells expressing GFP to cells expressing GFP-wildtype LS2. **(B)** PSI values for alternative exons that were significantly differentially included when comparing GFP samples to any LS2 sample were used to compare samples by principle component analysis (PCA). **(C)** For each LS2 sample, delta PSI (dPSI) values were calculated as the difference between PSI values in the LS2 sample and GFP. dPSI values for each mutant were then compared to dPSI values in wildtype LS2. **(D)** For each mutant and each alternative exon, the dPSI values in the mutant and wildtype LS2 were compared. Positive values indicate increased activity in the mutant relative to wildtype while negative values indicate decreased activity. **(E)** The locations of the G-rich motif GGNGGNG were calculated in the sequence surrounding alternative exons. Exons were divided into three classes, LS2-repressed, LS2-activated, and LS2-insensitive. For each class, as the sequence within and surrounding the exon is considered starting at nucleotide 0 (x-axis), the fraction of sequences in that class that contain a motif up to that point is calculated. This process is repeated as more and more sequence is considered. P values are Wilcoxon rank sum values comparing the cumulative density functions.

We then set out to compare the relative activities of wild type LS2 to mutant proteins that affect RNA binding *in vitro*, as shown above. To identify exons on which to make these comparisons, we identified a set of “LS2-sensitive” alternative exons that were significantly differentially included when comparing the GFP line to *any* LS2 line. To control for indirect effects stemming from the loss of free cellular dU2AF1 due to its titration away by exogenous LS2, we excluded from this set any exon that has been previously shown to be sensitive to a reduction in dU2AF1 levels^89^. A principal component analysis (PCA) of the PSI values of these 572 LS2-sensitive exons revealed that the GFP line and wild type LS2 line were well-separated, consistent with the strongest effect on splicing being observed in the wild type LS2 line **(Fig. 6B, Supplementary Fig. 12C**). The SF1_int mutant was essentially identical to wild type LS2 in this analysis with only modest reductions in activity, indicating that the interaction of LS2 with SF1 may be of little consequence for splicing regulation. The mutant showed modest reductions in activity. However, the U2AF1_int, RRM2, RRM1_beta5 and GS_linker mutants showed significant reductions in their splicing regulatory activity, as evidenced by their closer association with the GFP control samples in the PCA plot **(Fig. 6B, Supplementary Fig. 12C)**.

To further investigate the relative activity of wild type and mutant LS2, we calculated delta psi (ΔPSI) values for the LS2-sensitive exons by comparing PSI values in LS2 and control GFP lines **(Fig. 6C)**. We then directly compared the ΔPSI values of the LS2-sensitive exons in wild type and mutant LS2 lines. ΔPSI values showed a general correlation between wild type LS2 samples and all the mutants, indicating that exons were being similarly regulated by each LS2 mutant. However, the magnitude of ΔPSI was generally stronger in the wild type LS2 samples compared to the LS2 mutants, indicating that the mutants were generally weaker regulators of alternative splicing. As further evidence of this weakened activity observed in the mutants, we compared the magnitudes of ΔPSI values in LS2 wildtype and mutant lines **(Supplementary Fig 12D)**. We observed little to no effects on splicing activity in the SF1_int and GS_linker mutants but observed significantly weaker splicing activity in the U2AF1_int, RRM2, and RRM1_beta5 mutants, consistent with their reduced ability to bind RNA.

Through our structural and binding assays, we identified G-rich RNA sequences as prime targets for LS2 binding. To investigate if G-rich sequences mediate LS2 splicing regulatory activity, we asked if they were enriched near exons that show significant differential inclusion when comparing GFP and wild type LS2 lines. We found that exons whose inclusion was repressed by LS2 were significantly more likely to contain the G-rich motif GGNGGNG in their exonic or flanking intronic sequences than LS2-insensitive exons **(Fig. 6D**). Conversely, exons whose inclusion was activated by LS2 were less likely than LS2-insensitive exons to contain the motif. These findings are consistent with LS2 repressing exon inclusion through interaction with G-rich target sequences.

## Discussion

In this study, we have characterized structural and RNA binding properties of a tissue-specific alternative splicing factor LS2, which has evolved from the dU2AF2. Despite the high degree of sequence and structural homology with the single-stranded polypyrimidine tract binding U2AF2, we show that the LS2 RRMs recognize the G4 fold of the guanosine-rich RNA. Given the fundamental preference of canonical RRM domains towards the single-stranded nucleic acids^90^, our findings provide a novel insight to understand the versatile RNA-binding mode of RRM domains. RRMs have been reported to use various ways to mediate protein-RNA or protein-protein interactions depending on structural variations and extensions to the basic RRM fold^91-93^. Our structural and biophysical data show that both LS2 RRM domains have retained canonical βαββαβ fold including the consensus RNP2 and RNP1 motifs positioned at respective β1 and β3 strands. In comparison to U2AF2 we notice subtle conservative mutations in terms of RNA binding residues of LS2, that might explain the differential RNA binding by LS2. However, the N-terminal extension of the linker connecting the two RRMs is potentially crucial for its RNA-binding specificity that is discrete from its homologous U2AF proteins. We have shown these LS2-specific residues, composed of aromatic and positively charged residues, are not flexible and adopt a (partially) preformed α-helical conformation that weakly interacts with the α1 helix of the RRM2 domain. Importantly, this region is conserved throughout LS2 homologs from other *Drosophila* species. In the presence of G quadruplex RNA, this region exhibits significant chemical shift changes, which are likely the result of direct interaction with RNA nucleotides, although an indirect conformational effect cannot be ruled out. Previous studies have reported such transient helical regions in other nucleic binding proteins, where they become ordered upon binding to its target nucleic acid sequences ^90^ or modulate RNA binding selectivity ^44^. Although it is very intriguing to see that RNA-binding involves helix α_0_ in the N-terminal region of the linker, line broadening of NMR signals for the corresponding residues in the presence of RNA posed difficulties for further detailed analysis. The LS2 linker also contains a C-terminal region, which is predominantly composed of highly conserved hydrophobic residues among the U2AF2 paralogs. The linker-RRM2 solution NMR structure reveals that these C-terminal linker residues interact with β2 and β3 strands of RRM2, and thus gain partial rigidity as shown by NMR relaxation data. In the case of LS2, this interaction does not obviously alter the interaction of RRM2 with the G quadruplex RNA. However, the interaction may modulate the RNA binding by bringing the N-terminal α_0_ helix close to RRM2 and enable cooperative interactions of RRM2, the linker α_0_ helix and RRM1. Interestingly, LS2 RRM1,2 linker also contains five novel serine residues also in the α_0_ helical motif, which are highly conserved among the *Drosophila* LS2 homologs, and may serve as phosphorylation sites. Gene Ontology terms for the potential functions of LS2 also suggest its role in the gene regulation via phosphorylation ^45^. The LS2 phosphorylation could potentially regulate the aggregation-prone behavior of LS2, as observed in the case of elongation initiation factor 2α^94^.

We tested the ability of mutants designed based on our structural analysis to regulate alternative splicing in *Drosophila* S2 cells. Since LS2 is only very poorly expressed in S2 cells, our exogenously expressed wild type and mutant LS2 proteins comprised the vast majority of LS2 in the cells. Expression of wild type LS2 resulted in altered rates of inclusion for approximately 450 alternative exons. The majority of these exons were less included in transcripts upon LS2 expression, consistent with the previously reported role of LS2 as a splicing repressor ^45^. We found that mutations that destroyed the ability of LS2 to interact with the branch-point binding protein SF1 had little effect on the activity of LS2. However, those that interfered with the ability of LS2 to bind RNA resulted in less potent *in vivo* splicing regulatory activity. Importantly, the changes in exon inclusion exerted by these mutants tended to be of the same *sign* as was observed with wild type LS2 (i.e. increased or decreased exon inclusion) but were of smaller *magnitudes*. These observations are consistent with the RNA binding mutants acting as functional hypomorphs.

Our study shows that the LS2 protein recognizes G quadruplex (G4) RNA structures *in vitro*. We have also identified an enrichment of G-rich sequences near the vicinity of LS2 target exons. Various roles have implicated the regulation of RNA function by G quadruplex formation, highlighting a regulatory role of this local structure in the cellular context^95-99^. Recent studies show the presence of G4-forming motifs in the vicinity of the splice sites and also highlight their influence on splicing regulation ^60,73^ strongly suggesting that the formation of G4 structures could provide another degree of regulation in alternative pre-mRNA splicing. In fact, structure-forming, *cis*-regulatory elements including G4 can play an active role in the splicing by regulating the splicing reaction on their own without the need of splicing factors ^100,101^. The equilibrium between the single-stranded or the structured form of RNA can be the decisive factor for the recruitment of splicing factors depending upon their specificity for the sequence or the shape of RNA. Thus, G4 formation and its stability may have an impact on the ability of LS2 to regulate alternative splicing. A recently published study proposes the presence of the unfolded G4 structures in eukaryotic cells, proposing the presence of a robust G4 unfolding molecular machinery^102^. However, G4 formation has also been reported to be cell cycle and cell state dependent ^64,75,103^. Thus, cellular G4-unfolding activity may itself be subject to regulation. Moreover, the use of steady-state measurements might as well fail to detect transient G4 formation^104-106^.

Due to the poor quality of recombinant LS2 protein and its RNA complex high-resolution structural studies by using X-ray crystallography and NMR, information of the novel RRM1,2-G4 interaction remain elusive. Although there is still very limited structural information available regarding the G4 recognition by proteins, recently reported structures of bovine DHX46 bound to ssDNA-G4^107^ and the Rap1 DNA binding domain in domain in complex with DNA G4 ^108^ insight into the recognition mode. It is important to note that both structures highlight that specificity for the G-quadruplex recognition is provided by an α-helix (composed of basic and hydrophobic residues) covering the topmost planar face of the G4 structure as well as salient contributions from arginine side chains hydrogen bonding to the phosphate backbone. The importance of arginine residues for G4 recognition was also observed from the structure of RGG peptide of FMRP bound to the RNA duplex-quadruplex junction ^109,110^. These findings are perfectly in line with our studies and strongly support our observation that the unique α_0_ helix in the LS2 RRM1,2 linker, which also harbors two arginine residues along with two aromatic and hydrophobic residues might be crucial in the G4 recognition by LS2. Moreover, the conservative mutations in LS2 RRM2 domain in comparison to U2AF2 such as K311R, K312R might provide the crucial hydrogen bonding along with the differential base stacking patterns mediated by the changes in the key aromatics (F245Y and Y285F) of the RNP sites.

In conclusion, our detailed structural, biophysical and biochemical characterization of the LS2 RRM1,2 RNA binding region shows that LS2 recognizes the G4-fold of the guanosine-rich target RNA, distinct from the common recognition of single-stranded RNAs by homologous RRM domains. The recognition of structured *cis*-regulatory RNA motifs likely plays important roles in the regulation of alternative splicing and has been exploited in the evolution with the U2AF2 paralog LS2 for tissue-specific splicing regulation.

## Online Methods

### Cloning, expression, and purification

LS2 constructs RRM1 (110–204), RRM2 (242–319), linker-RRM2 (204–325), RRM1,2 (110–325), RRM1,2 Δlinker (110–319 without 205–240) were subcloned into pET24d-modified vectors by using appropriate primers with NcoI and XhoI restriction sites. In the case of RRM1, RRM2 and RRM1,2 Δlinker maltose binding protein and thioredoxin tags were used as fusion tags, respectively, in addition to an N-terminal hexahistidine tag and TEV cleavage site. For RRM1,2 and linker-RRM2, sumo fusion tag was used along with N-terminal hexahistidine tag and a sumo protease cleavage site. Recombinant protein constructs were transformed and expressed in *E. coli* BL21 (DE3) cells using either Luria-Bertani (LB) or minimal M9 media supplemented with ^15^N NH4Cl or ^15^N NH4Cl and ^13^C-glucose. Protein expression was induced by the addition of 1 mM isopropyl β-D-thiogalactoside (IPTG) at an OD600 nm of 0.6. Following induction, bacterial cultures with RRM1 and RRM2 constructs were incubated at 16 °C whereas cultures with RRM1,2; RRM1,2 Δlinker and linker-RRM2 were incubated at 37 °C for 16 hours on a rotary shaker (220 rpm). Bacterial cells were harvested by centrifugation at 4000 rpm for 20 min at 4 °C. The cell pellet was resuspended in the lysis buffer consisting of 20 mM Tris, pH 8.0, 500 mM NaCl, 5 mM imidazole, 1mM β-mercaptoethanol supplemented with AEBSF-HCl inhibitor, DNaseI (1 mg/ml) and MgSO4 (10 mM). Cells were lysed by sonication on ice followed by the centrifugation at 19000 rpm for 45 min at 4 °C.

In the case of the RRM1 and RRM2 constructs, cleared lysate was loaded onto the pre-equilibrated Ni^+2^-NTA resin (Qiagen). Further, washing with 10 column volumes of the lysis buffer supplemented with 25 mM imidazole was performed to remove non-specifically bound proteins. Bound protein was eluted by applying the elution buffer consisted of lysis buffer with 250 mM Imidazole. Overnight dialysis and cleavage using TEV protease in 10 mM Tris, pH 8.0, 150 mM NaCl, 1 mM β-mercaptoethanol was carried out. Second IMAC purification was performed to separate cleaved solubility tag, TEV protease and uncleaved protein. Amicon ultra-15 (Millipore) concentrator was used to concentrate the flow-through consisting of the target protein up to 1-5 ml. It was then applied onto HiLoad 16/60 Superdex 75 (GE Healthcare Biosciences) size-exclusion column in 10 mM Tris, pH 8.0, 150 mM NaCl, 5 mM β-mercaptoethanol buffer for further purification. Fractions containing pure protein were collected and buffer exchanged to 20 mM potassium phosphate, pH 6.5, 50 mM NaCl, 5 mM DTT using the Amicon concentrator for further experiments.

For the linker-RRM2 and RRM1,2 following the sonication of cells and subsequent centrifugation, pellets containing inclusion bodies were used for the subsequent purification steps. Pellets were dissolved in the fresh buffer containing 20 mM Tris, pH 8.0, 100 mM NaCl, 5 mM imidazole, 1mM β-mercaptoethanol and 8 M urea, by performing extensive sonication on the ice. Centrifugation at 19000 rpm for 30 min at 4 °C was implemented to remove the undissolved inclusion bodies. The supernatant was applied onto the preequilibrated Ni^+2^-NTA resin and subsequent washing and elution steps were performed as mentioned above but in the presence of 8 M urea. Next, overnight dialysis in 10 mM Tris, pH 8.0, 150 mM NaCl, 1 mM β-mercaptoethanol buffer was performed followed by cleavage using yeast sumo protease (dtUD1). Subsequently, second IMAC purification and gel filtration steps were performed as mentioned above.

Sodium dodecyl sulfate polyacrylamide gel (SDS-PAGE) in combination with Coomassie-blue staining was performed to check the purity of the protein samples in each step. Concentration of the pure protein fragments was measured by absorbance at 280 nm, using Nanodrop 1000 spectrophotometer (Thermo Scientific).

Site-specific LS2 mutants were prepared with the Quick-Change site-directed mutagenesis kit (Stratagene) and mutations were verified by the DNA sequencing.

### RNA samples

Unlabeled RNA oligonucleotides 21-mer 5’-(GGGGAGGAGGGGGGCGUAUGA), 14mer 5’-(GGGUGGUGGAGGGG), 8mer 5’-(GGUGGUGG), 7mer 5’-(CGUAUGA) were purchased from IBA GmbH, Göttingen (Germany) in lyophilized form. Oligonucleotides were dissolved in sterile water, followed by heating at 95 °C for 3 mins and snap cooling on ice or RT for subsequent use. Concentrations were determined with a NanoDrop 1000 spectrophotometer (Thermo Scientific) at 254 nm. For NMR experiments, RNA samples were dissolved in 90% H_2_O/10% D_2_O or 100% D2O to a final volume of 180 μL (for 3mM NMR tubes) or 300 μL (for Shigemi tubes) with concentrations range of 30–500 μM.

### NMR spectroscopy

NMR spectra were recorded on an AVIII500, AVIII600, AVIII750, AVIII800, AVI900 and an AVIII950 spectrometer, equipped with cryogenic or room temperature (for 750 MHz) triple resonance gradient probes. For data acquisition, 50–1000 μM protein samples were used in 20 mM potassium phosphate buffer, pH 6.5, 50–300 mM NaCl, 5 mM DTT along with 10% D_2_O added for the lock. Protein backbone resonance assignments were derived from experiments such as HNCA, HNCACB, CBCA(CO)NH^111^. Amino acid side chain resonances were assigned from HCCH-TOCSY, (H)CC(CO)NH, ^15^N- and ^13^C-edited NOESY-HSQC experiments. (HB)CB(CGCD)HD, (HB)CB(CGCDCE)HE^112^, ^1^H-^13^C HSQC, as well as Aromatic ^13^C-edited NOESY-HSQC, were used for assignments of the aromatic resonances. Stereospecific assignments were performed by the method described by^113^. Data processing was performed by using NMRPipe/Draw^114^ or Topspin 3.2 (Bruker) and analysis was performed by using NMRFAM-SPARKY^115^.

Chemical shift derived secondary structure was determined using random coil chemical shifts derived using POTENCI algorithm via webserver(https://st-protein02.chem.au.dk/potenci/)^116^. To highlight regular secondary structural elements, the obtained raw data were treated with 1-2-1 smoothening function for residues (*i*-1)-(*i*)-(*i*+1) as described before^117^. Backbone ^15^N relaxation data were acquired with 250 μM protein at 298 K on a 500 MHz Bruker NMR spectrometer. *R*_1_, *R*_1ρ_, and steady-state heteronuclear {^1^H}-^15^N-NOE experiments were performed as described^118,119^. Ten different relaxation delay points together with two redundant delays such as 4/4, 8, 16, 32, 48, 64/64, 84, 96, 120, 160 ms were recorded for *R*_1_ data measurement. *R*_1ρ_ data measurement was performed by recording 8 different relaxation delay points with two redundant delay points such as 5/5, 10, 20, 40, 65/65, 90, 120, 160 ms. Duplicate time points were taken into account for error estimation. *R*_2_ relaxation rate was derived from each residue from calculated *R*_1ρ_ relaxation rate using the relation *R*_1ρ_ = *R*_1_ cos^2^θ + *R*_2_ sin^2^θ, where θ = tan^−1^ (ν_1_/Δν) and Δν is the offset of the rf field to the resonance. {^1^H}-^15^N heteronuclear NOE spectra were recorded with and without ^1^H saturation of 3 s Error bars for the heteronuclear NOE data are calculated by error propagation of peak height uncertainties using average noise levels in the NMR spectra. All relaxation experiments were recorded as pseudo-3D experiments. NMRFAM-SPARKY ^115^ was used for analyzing the relaxation data and for the calculation of relaxation rates along with the error determination.

### Structure calculation

CYANA 3.0 with automated NOESY assignment ^120^ was used to generate solution structures of the RRM2 and linker-RRM2. TALOS+^121^ was used to generate *φ* and *ψ* backbone dihedral angle restraints based on chemical shifts. These angle restraints along with distance restraints derived from NOEs were used for restrained molecular dynamics simulations and simulated annealing during structure calculation protocol. Water refinement was performed by CNS/ARIA^122^. All structures were validated using wwPDB Validation Service (validate-rcsb.wwpdb.org) whereas PyMol (Schrödinger) was used to obtain the molecular images. The structural statistics for linker-RRM2 and RRM2 are reported in Table 1.

**Table 1.** Structural statistics of the LS2 Linker-RRM2 and RRM2 NMR structures.

|  | Linker-RRM2 | RRM2 |
| --- | --- | --- |
| **Completeness of $^1\text{H}$ chemical shift assignments (%)** | 91.6 | 96.3 |
| **NMR restraints** |  |  |
| **NOE-derived distance restraints** | 1691 | 1724 |
| short-range, $\vert i-j\vert \leq 1$ | 895 | 783 |
| medium-range, $1 < \vert i-j\vert < 5$ | 280 | 314 |
| long-range, $\vert i-j\vert \geq 5$ | 516 | 627 |
| **Dihedral restraints ($\phi/\psi$ from TALOS)** | 146 | 116 |
| **Restraints violations** |  |  |
| rms distance violation ($\text{\AA}$) | 0.0105±0.0010 | 0.0082±0.0006 |
| max. distance violation ($\text{\AA}$) | 0.31±0.05 | 0.27±0.03 |
| rms dihedral violation ($^\circ$) | 0.811±0.043 | 0.454±0.045 |
| max. dihedral violation ($^\circ$) | 5.06±0.43 | 3.05±0.36 |
| **RMSD to mean structure statistics ($\text{\AA}$)** |  |  |
| backbone atoms | 0.28 | 0.26 |
| heavy atoms | 0.76 | 0.70 |
| **Cyana target function value** |  |  |
| First cycle | 270.44 | 82.76 |
| Final | 3.61 | 1.51 |
| **Ramachandran plot statistics ($\text{\AA}$)<sup>a,b</sup>** |  |  |
| Residues in most allowed regions (%) | 90.9 (84.8) | 93.8 |
| Residues in additionally allowed regions (%) | 7.7 (13.1) | 5.6 |
| Residues in disallowed regions (%) | 1.3 (2.1) | 0.5 |

<sup>a</sup> calculated using RAMPAGE<sup>126</sup> for the 10 lowest energy structures after water refinement
<sup>b</sup> for the structured region (242-318). Values in the parentheses correspond to the total residues of linker-RRM2 structures.

To determine the structure of the RRM1 domain CS-ROSETTA structure calculation approach^123^ was used. 10,000 all atom models were generated. In addition, 25 strong, long range (NOE) distance restraints were included to guide the process. Structural statistics for RRM1 are reported in Table 2.

**Table 2.** Structural statistics for the LS2 RRM1 CS-ROSETTA structure.

|  |  |
| --- | --- |
| **NOE-derived distance restraints** |  |
| long-range, $\vert i-j\vert \geq 5$ | 25 |
| **Cordinate precision RMSD (Å)** |  |
| backbone atoms | 0.28 |
| heavy atoms | 0.76 |
| **Ramachandran plot statistics (Å)<sup>a</sup>** |  |
| Residues in most allowed regions | 98.6% |
| Residues in additionally allowed regions | 1.4% |
| Residues in disallowed regions | 0.0% |

<sup>a</sup> calculated using RAMPAGE<sup>126</sup> for the 10 final structures with lowest energy and ROSETTA score

### SAXS

SAXS data were recorded on the in-house Rigaku BioSAXS1000 instrument equipped with the Rigaku HF007 microfocus rotating anode with a copper target (40 kV, 30 mA). Measurements were carried out in 20 mM potassium phosphate buffer, pH 6.5, 300 mM NaCl, 5 mM DTT at 277 K. 1,2,3 mg/ml concentrations of RRM1,2 and RRM1,2 Δ linker protein samples, as well as buffer alone for solvent subtraction, were used for measurements. Sample measurement was performed in four frames with 900 s exposure time per frame. Frames were compared to check for the radiation damage, averaged and solvent subtracted by the SAXSLab software (v3.0.2). Data was processed using the ATSAS software suite^124^.

### NMR titration experiments

^1^H-^15^N HSQC or ^1^H-^15^N SOFAST-HMQC^125^ REF spectra were used to monitor the protein backbone amide chemical shifts whereas RNA imino resonances were tracked by 1D water-flip-back WATERGATE experiments each step of titration. Experiments were performed in the 20 mM potassium phosphate buffer, pH 6.5, 50 mM NaCl, 5 mM DTT at 298 K, unless stated otherwise. 50-100 μM ^15^N or ^15^N-^13^C protein sample was titrated with increasing amounts of RNA oligonucleotides typically with the steps of 0, 0.1, 0.2, 0.4 upto 1.0, 2.0 or 4.0 molar equivalents of RNA to protein. Chemical shift perturbations were calculated using the relation 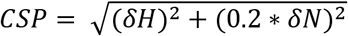, where *δH* is the difference in the proton chemical shift in parts per million (ppm) and *δN* is the difference in the nitrogen chemical shift in ppm.

### Isothermal titration calorimetry

For ITC studies PEAQ-ITC (Malvern Instruments) was used. Protein samples were dialyzed extensively in the buffer used for the titration. RNA oligonucleotides were also solubilized in the same buffer to provide the baseline as required. 20 μM 21-mer RNA was used, in which 200 μM RRM2/linker-RRM2 was titrated. 19 injections comprised of 2 μL titrant were performed with 150 s spacing in between each injection. Experiments were performed at 7 °C with stirring at 750 revolutions per minute (rpm). Data processing was performed by the ITC Analysis software provided by the Malvern Instruments.

### Circular dichroism

The CD spectroscopy was performed using Jasco J-810 spectrometer using 1 mm path length cuvette (Hellma Analytics). The experiments were conducted using the wavelength scan of 200-350 nm. Each spectrum was recorded with 3 or 10 number of scans, 1 s response time and a 2 nm bandwidth. Experiments were performed on 40 μM or 60 μM sample with 300 μL volume at 298 K.

### Static light scattering (SLS)

Superdex 75 16/60 size-exclusion column (GE Healthcare) was used to perform the SLS experiments. Refractive index (RI) and light scattering by the sample components were detected by Viscotek Triple Detector Array (Malvern instruments). 100 μL of sample volume was injected on to the column, which was pre-equilibrated with the buffer. The flow rate of 0.5 ml/ml was used for each run. Calibration was performed by using 4 mg/ml BSA standard. For data analysis, OmniSEC software was used.

### Expression of LS2 mutants in S2 cells

Wild type and mutant LS2 constructs were expressed as GFP fusions (LS2 N-terminus, GFP C-terminus) in pUChygMT. This plasmid contains a selectable hygromycin resistance gene as well as a copper-inducible metallothionein promoter. Stable S2 cell lines were constructed expressing the LS2 constructs by transfecting the cells with the constructs using Effectene (Qiagen). Transfections were performed in one well of a six-well plate using 2μg plasmid DNA, 16μL enhancer, 20μL Effectene, and 150μL EC buffer. 48 hr after transfection, selection was started using 100μg/mL hygromycin. The cells were selected for approximately 4 weeks in 100 μg/mL hygromycin.

To induce expression of LS2, the cells were treated with 500μM copper sulfate for 53 hr. Equal expression of LS2 among the GFP control, wild type and mutant cell lines was verified by immunoblotting using an anti-GFP antibody (Abcam ab290). The cells were then lysed and RNA was collected using the Qiagen RNeasy Mini kit. RNA quality was verified using an Agilent Bioanalyzer, and stranded, paired-end libraries were prepared using 25 ng total RNA on the Illumina NeoPrep liquid handler.

For imaging of the subcellular localization of LS2, cells were plated on #1.5 PLL-coated coverslips (Neuvitro). After induction with copper for 72 hr, the coverslips were mounted on slides and imaged at 60X on a Deltavision Elite widefield deconvolution microscope (GE). Images were collected in the FITC (GFP-LS2) and DAPI (DNA) channels.

### Analysis of mutant LS2 splicing activity

Libraries were sequenced on an Illumina NextSeq sequencer using 2×75 paired-end sequencing. Approximately 40 million paired-end reads were obtained per sample. Reads were mapped against the Drosophila dm6 genome annotation using STAR (Dobin et al, Bioinformatics, 2013). The genome annotation was downloaded from Ensembl, and was BDGP6, Ensembl version 88. For gene expression analysis, samples were quantified using kallisto (Bray et al, Nature Biotechnology, 2016). For splicing analysis, samples were analyzed using rMATS (Shen et al, PNAS, 2014). Exons were designated as significantly affected in a GFP control versus LS2 sample if the FDR for the exon was less than 0.05 and the absolute value of the difference in PSI values between the samples was greater than or equal to 0.05. PSI value comparisons and motif analyses were done with custom Python and R scripts.

For the identification of exons whose inclusion was sensitive to dU2AF1 depletion, RNAseq data from S2 cells treated with siRNA against dU2AF1 was downloaded from the Short Read Archive at SRP0556965. The reads were aligned and quantified as above for the LS2 expression experiments. Quantification of exon inclusion in these samples was also performed as above.

To quantify motif occurrences near LS2-regulated exons, sequences surrounding these regulated exons were extracted (see **Fig. 6D**). 50 nucleotides of exonic sequence and 200 nucleotides of intronic sequence were considered for each regulated alternative exon as well as its flanking exons. If the size of the flanking intron was less than 200nt resulting in this sequence bleeding into the next exon, the sequence was truncated at the exon/intron boundary. Sequences were analyzed beginning in the exonic sequence and walking toward in the intronic sequence. For each sequence at each position, the existence of a motif of interest, in this case GGNGGNG, was queried. The position of this motif within the walk is then recorded. The abundance of motifs is then plotted as a cumulative density function (CDF) along the sequences analyzed. The statistical significance of motif enrichments or depletions was calculated using a Wilcoxon rank sum test comparing the CDF of LS2-regulated exons to control, unaffected exons.

Raw sequencing data is available at GEO, https://www.ncbi.nlm.nih.gov/geo/, accession code GSE156213.

## Acknowledgements

We thank Gerd Gemmecker and Sam Asami for help with NMR experiments, Alexander Beribisky, Johannes Günther and Hamed Kooshapur for helpful discussions. We acknowledge SAXS measurements at the facility of the SFB1035 at Department Chemie, Technical University of Munich. We acknowledge the Deutsche Forschungsgemeinschaft (DFG) grants GRK1721 and SFB1035 (to M.S.), and US NIH grants R35GM118121 and R21HG010238 (to D.C.R.) for financial support.

## Author contributions

A.A.K. prepared protein samples for biochemical and structural biology, and performed structure calculations; J.M.T. performed *in vivo* splicing assays and analyzed RNA sequencing data; H.S.K. provided project directions and helped with NMR data analysis; C.H. performed CS-ROSETTA calculations; A.G. and R.S. performed SLS and SAXS data measurements, respectively; D.C.R. and C.B. designed *in vivo* splicing assays and RNA sequencing experiments. M.S., J.M.T. and D.C.R conceived the project; A.A.K., J.M.T., D.C.R. and M.S. wrote the manuscript; All authors discussed the project and approved the manuscript.

