## Supporting Information for "Structural evolution of the tissue-specific U2AF2 paralog and alternative splicing factor LS2"

### Supplementary results

**Isolated LS2 RNA binding RRM domains adopt a canonical RRM fold in solution.**

We observe that chemical shift derived secondary structures of both isolated LS2 RRM domain are fully consistent with the CS-ROSETTA model of LS2 RRM1 and solution NMR structure of LS2 RRM2. $^{15}$N relaxation data show that both domains are well folded and are mostly rigid with some increased flexibility in loops (**Supplementary Fig. 2**). A multiple sequence alignment shows that loop2 between the α1 helix and a β2 strand of LS2 RRM1 (143–147) is longer in LS2 compared to U2AF2 orthologs (**Supplementary Fig. 1)** and also contains residues such as K143, H146 and Y147 which could mediate RNA binding. FUS RRM is reported to contain longer connecting loop (loop2) between α1 helix and a β2 strand, which mediates RNA interactions[1]. NMR secondary chemical shifts and the relaxation data show that residues in this region exhibit higher flexibility and thus lack any secondary structure formation or transient interaction with other residues. NMR titration data reveals that in contrast to FUS, LS2 RRM1 loop2 does not participate in RNA binding **(Supplementary Fig. 7C)**.

**Imino resonance assignments and G4 fold of the 21-mer G4 RNA**

To confirm the G4 topology of 21-mer RNA in the presence of low $K^+$ concentration we employed two dimensional (2D) NMR spectroscopy. The resonances are tentatively labelled as 1–8 depending upon their position in the $^1$H NMR spectrum. Lower temperature (278K) was chosen to reduce the solvent exchange rate of the imino resonances, where only eight imino resonances are observable with 4 overlapped duplets (namely resonance 3, 4, 6, 7) **(Fig 2A)**. $^1$H-$^1$H NOESY in the presence of 5 mM KCl in $H_2O$ at 278K resulted in the highly overlapping imino cross peaks. Hence, we recorded imino NOESY spectrum in $D_2O$, which resulted in the fewer cross peaks and aided the resonance assignments of the core region of the G4, as well as two more imino resonances partially protected from the solvent exchange **(Supplementary Fig. 6A)**. The imino resonances showing high resistance for solvent exchange (namely resonances 1, 2, 3 and 5) might stem from the central plane of the G4 structure as seen by cross peaks for 1-2, 1-5, 3-2, 3-5. The NOE cross peaks between resonance 8-2 and 6-5 are consistent with the existence of the hexad structure formation, where imino 8 and 6 shows partial resistance for the solvent exchange despite being exposed to the solvent. After finding the correlations between the central plane of the G4 structure, imino NOESY in water was used to correlate the cross peaks between remaining resonances **(Supplementary Fig. 6B)**. The top plane is formed by resonances 8, 6, 4 and 3 as seen by the cross peaks between (8-4, 8-3, 6-4, 6-3) and bottom plane (4, 7, 6, 7) as seen by the cross peaks between (4-6,4-7, 7-6). Imino cross peaks between the planes such as (2-8, 2-4, 1-4, 1-7,3-4, 3-6, 5-6., 5-7) are consistent with the G4 structure with three planar topology.

Although, the 21-mer RNA sequence does not contain the continuous stretch of GGGAAGGG motif characteristic of G4 formation, interspersed guanosine nucleotides may correspond to the connecting loops adjoining G-quartets thus, forming the proposed G4 structure with three planar topology where varying length of guanosine loop residues contribute to the distinct homodimeric structure **(Supplementary Fig. 6C)**.

**LS2 RRM2 shows specificity towards 21-mer specific G4 structure**

In order to further assess the specificity of the LS2 RRMs towards the G4 RNA structure, we performed titrations of the individual RRM domains with shorter guanosine-rich oligonucleotides namely 14-mer (GGGUGGUGGAGGGG), 8-mer (GGUGGUGG) and 7-mer (CGUAUGA). By screening various salt conditions, we observe that 14-mer predominantly shows a heterogeneous G4 population irrespective of the $K^+$ concentration (**Supplementary Fig. 9A**). In contrast, similar to 21-mer RNA, 8-mer guanosine-rich RNA adopts homogeneous G4 at low $K^+$ concentration (**Supplementary Fig.9B**). NMR titration experiments of RRM1 with the 14-mer and 8-mer RNAs show small CSP that map to similar RNA binding interface as found with the 21-mer RNA (**Supplementary Fig. 9C, 9E, 10A**). These data suggest that RRM1 interacts non-specifically with the RNA. NMR titration of RRM2 with the 14-mer RNA show no significant spectral changes, while in the presence of 8-mer RNA minor CSP changes are observed that are much smaller than those seen for the 21-mer RNA (**Supplementary Fig. 9D, 9F, 10B**). It is important to note that 14-mer contains all the nucleotides present in the 8-mer RNA but even at low $K^+$ concentration, it adopts multiple conformations as opposed to the formation of uniform conformation by the 21-mer and the 8-mer RNA. Thus, suggesting G4 structure specific interaction by LS2 RRM2. Extensive line broadening of resonances is observed for both RRM and RNA resonances, which indicate the binding in multiple registers and/or in the intermediate exchange regime. The weak interaction of RRM2 with 8-mer RNA may result from a different G4 structure lacking hexad structure. Together these data indicate that RRM2 provides the specificity towards the homogeneous G4 conformation adopted by the 21-mer RNA in the presence of low $K^+$ concentration. To rule out the possibility that RRM2 shows specificity for RNA sequence elements in the 21-mer RNA, which are not found in the 14-mer or 8-mer RNA, we performed a titration of the RRM2 with a non-G sequence present at 3' end of the 21-mer RNA i.e. 7-mer RNA (CGUAUGA). No significant spectral changes are observed for the RRM2 amide signals upon addition of this RNA, indicating that RRM2 does not recognize this sequence (**Supplementary Fig. 10C**). Next, to check whether RRM2 interacts with free guanine nucleotides, which lacks the ability to adopt the G4 structure, the titration of GTP with RRM2 was performed. The addition of GTP also didn't have any significant effect on the RRM2 resonances as monitored by the $^{1}H$-$^{15}N$ HSQC (**Supplementary Fig. 10D**). This data indicates that RRM2 does not interact with free GTP nucleotides either and thus, confirms its specificity towards the uniform G4 structure adopted by the 21-mer RNA.

#### Supplementary Figures

##### Supplementary Figure 1

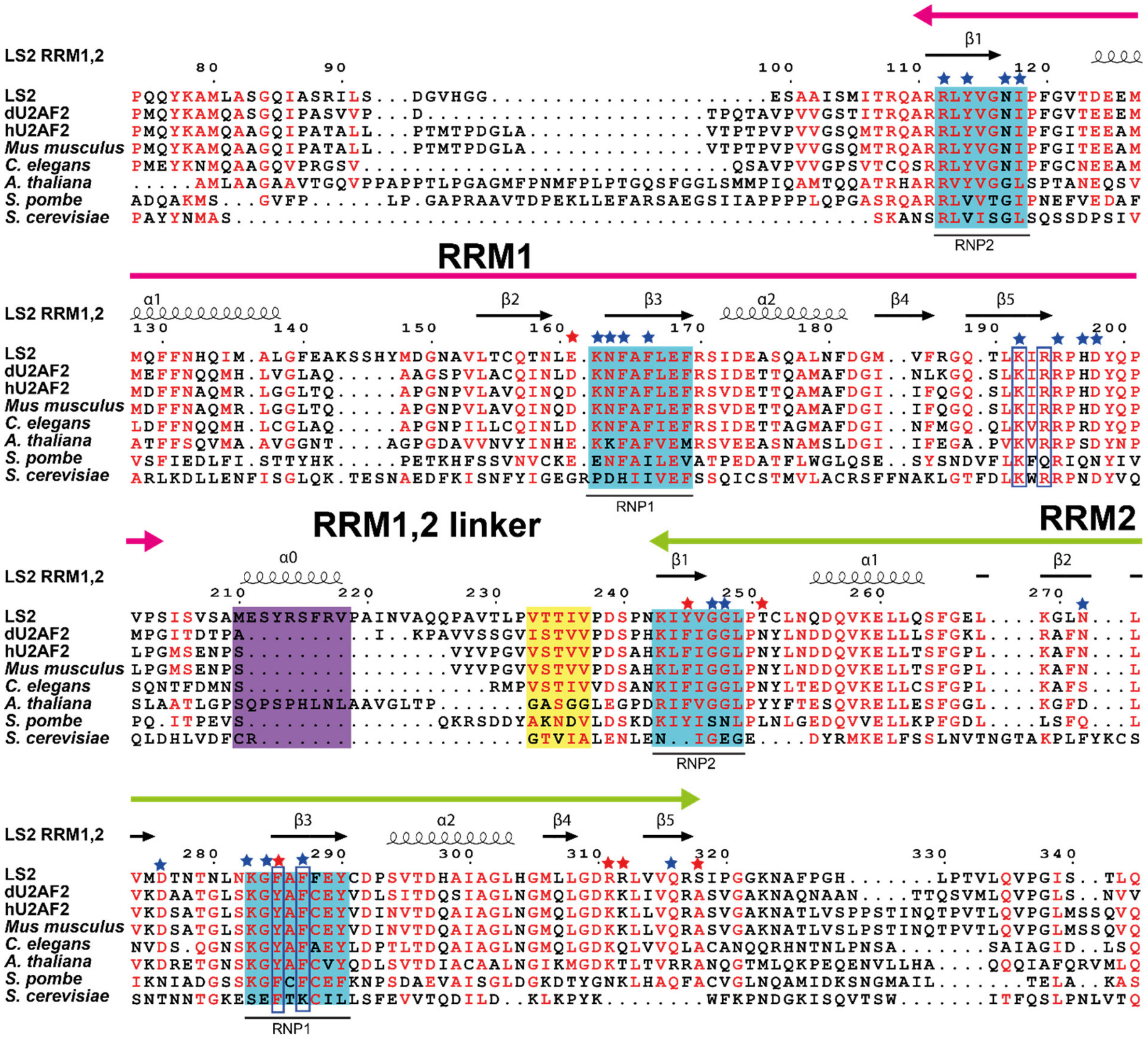


**Supplementary Figure 1. Multiple sequence alignment of LS2 with U2AF2 from different eukaryotic species.** Original sequences used in the analysis [residues (74–343) of *Drosophila melanogaster* LS2, residues (56–311) of *Drosophila melanogaster* dU2AF2, residues (104–369) of *Homo sapiens* hU2AF2, residues (104–369) of *Mus musculus* U2AF2, residues (126–366) of *Caenorhabditis elegans* U2AF2, residues (182–465) of *Arabidopsis thaliana* U2AF2, residues (147–411) of *Saccharomyces pombe* U2AF2, residues (195–415) of *Saccharomyces cerevisiae* U2AF2 were aligned by Clustal Omega[2] and analyzed using ESPript 3.0[3]. Secondary structural elements indicated on the top of the sequence alignment corresponds to the CS-ROSETTA model of LS2 RRM1 and solution NMR structure of LS2 RRM2. The LS2-specific linker residues with α-helical propensity are highlighted with a purple background whereas RRM2 interacting linker residues are highlighted with a yellow background. The RNP1 and RNP2 sequence motifs are colored with cyan background. Violet boxes indicate the residues used for mutational analysis from LS2 RRM1 and RRM2, respectively. Blue and red stars mapped on top indicate polypyrimidine tract binding hU2AF2 residues[4] which are conserved and not conserved in LS2 protein sequence, respectively.

#### Supplementary Figure 2

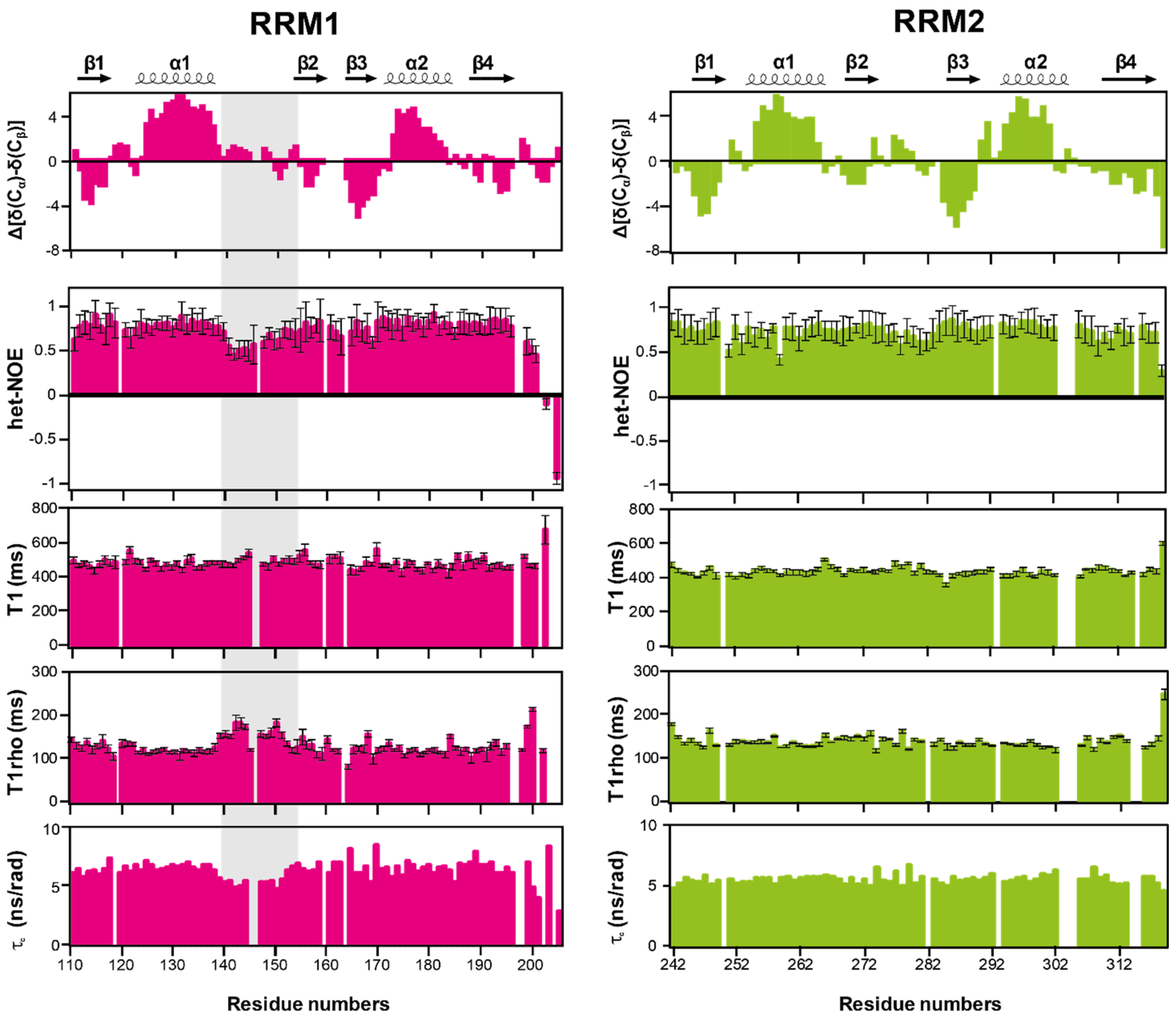


**Supplementary Figure 2. NMR secondary chemical shifts, secondary structure and $^{15}$N relaxation of isolated LS2 RRM domains.** Chemical shift derived secondary structure and $^{15}$N NMR relaxation data for the LS2 RRM1 (left panel) and RRM2 (right panel) in the free form. The secondary structure elements are shown on top. LS2 RRM1 specific residues of the loop2 are highlighted in a grey background.

#### Supplementary Figure 3

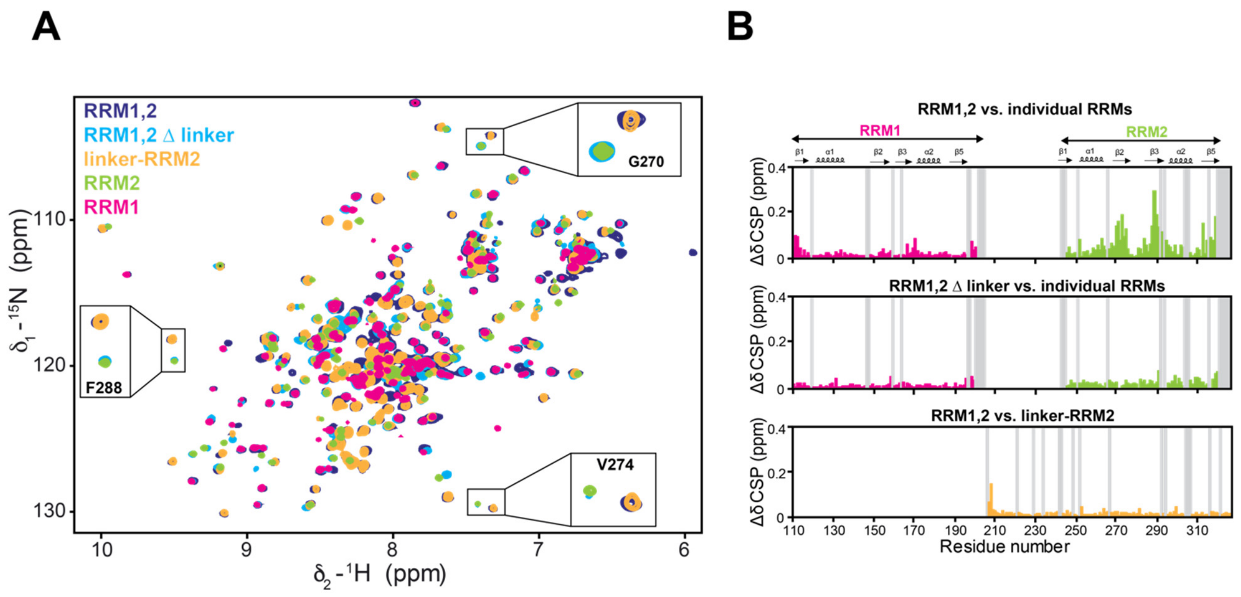


**Supplementary Figure 3. LS2 RRM1-RRM2 interdomain linker mediates interaction with RRM2. (A)** The overlay of the $^{1}$H-$^{15}$N HSQC of various RNA binding domain fragments of LS2 recorded in 20 mM potassium phosphate buffer pH 6.5, 300 mM NaCl, 5 mM DTT at 288 K. Selected residues of RRM2 exhibiting differential NMR chemical shifts are shown in insets. **(B)** The CSP analysis of the RRM1,2; RRM1,2 Δlinker with respect to individual RRM domains (top and middle panel, respectively), and RRM1,2 with respect to linker-RRM2 protein fragment (bottom panel). Grey bars indicate either missing assignments, proline residues or terminal residues. Residues undergoing chemical shift changes in the presence of the linker are located on the β2- and the β3-strands of the RRM2 whereas no significant chemical shift changes are observed in RRM1,2 in comparison to linker-RRM2 construct.

#### Supplementary Figure 4

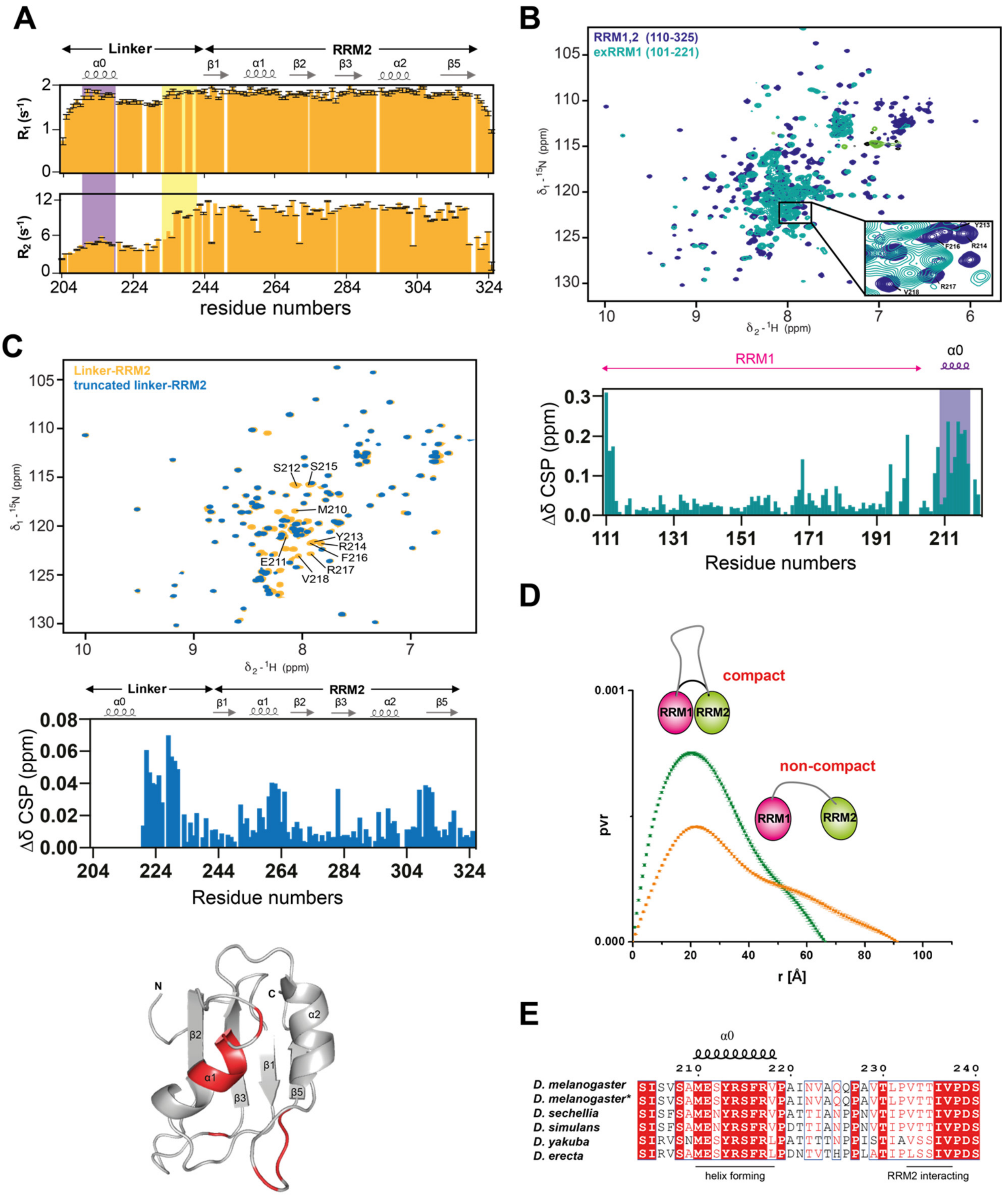

**Supplementary Figure 4. Analysis of LS2 RRM1-RRM2 interdomain linker. (A)** $^{15}N$ $R_1$ and $R_2$ relaxation rates of the linker-RRM2 highlighted with two semi-rigid linker regions corresponding to the linker-helix and

RRM2 interaction in a purple and a yellow background, respectively. **(B)** The overlay of the $^{1}H$-$^{15}N$ HSQCs (top panel) and CSP plot (bottom panel) of the RRM1,2 and exRRM1 constructs showing differential chemical shifts in the linker helix region. Data was acquired in 20 mM potassium phosphate buffer pH 6.5, 300 mM NaCl, 5 mM DTT at 288 K **(C)** The overlay of the $^{1}H$-$^{15}N$ HSQCs of the linker-RRM2 and truncated linker-RRM2 lacking the N-terminal of the linker (204–220), which include the helix forming residues in the LS2 RRM1,2 linker (top panel), CSP plot (middle panel) and residues undergoing CSPs higher than twice the standard deviation are plotted in red onto the structure of linker-RRM2 (bottom panel). Spectra were recorded in 20 mM potassium phosphate buffer pH 6.5, 300 mM NaCl, 5 mM DTT at 288 K. **(D)** SAXS analysis showing pair-distance distribution functions for RRM1,2 (orange) and RRM1,2 Δlinker (green). Both LS2 RRM1,2 and RRM1,2 Δ linker exist as a dumbbell shape in solution. RRM1,2 shows an ensemble of conformations with compact and non-compact states, as seen by the presence of the two lobes, whereas RRM1,2 Δ linker shows the existence of the compact state predominantly. **(E)** Multiple sequence alignment of the LS2 homologs from various *Drosophila* species. Residues used for sequence analysis are as following: residues 204–240 of *D. melanogaster*, residues 204–240 of *D. melanogaster*, residues 200–236 of *D. sechellia*, residues 200-236 of *D. simulans*, residues 193–229 of *D. yakuba*, residues 196–232 of *D. erecta*. Residues showing α-helical propensity as well as RRM2 interaction are marked.

#### Supplementary Figure 5

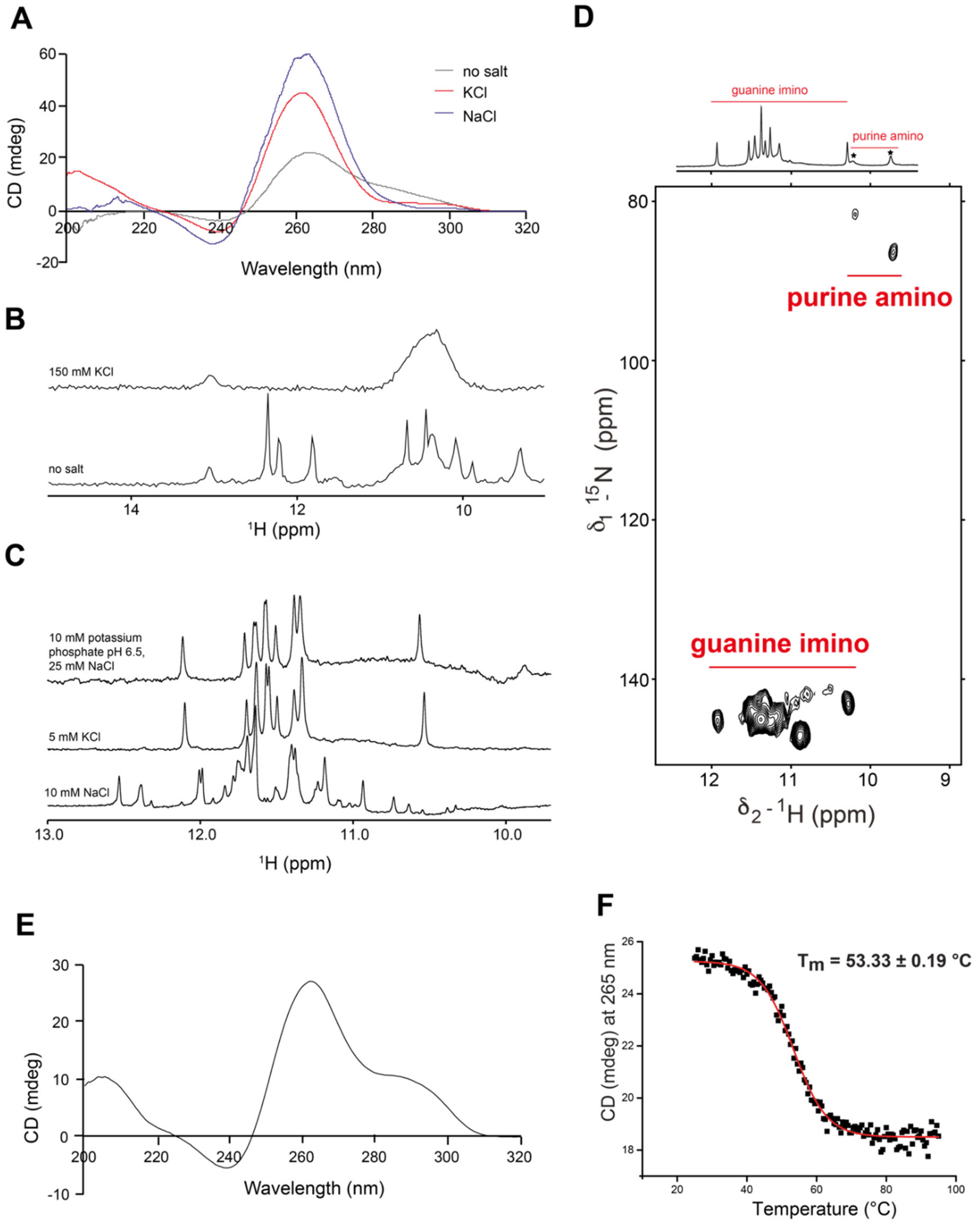


**Supplementary Figure 5. Biophysical characterization of G4 fold adopted by LS2 target poly G RNA sequence (21 mer) in the presence of various salt types. (A)** Circular Dichroism spectra of 50 µM 21-mer RNA in the absence of salt, in the presence of 160 mM NaCl and 160 mM KCl acquired at 25°C. A negative CD absorption at 240 nm and a positive absorption at 265 nm indicating the formation of a parallel-stranded G4 structure with more stability in the presence of salt. **(B)** $^1$H 1D NMR spectra of the imino region of 21-

mer RNA at in the presence of 150 mM NaCl and in the absence of salt recorded at 278 K. Imino resonances observed in the wobble base-pairing region (10-12 ppm) indicate G4 formation. **(C)** $^1$H 1D NMR spectra of the imino region of 21-mer RNA at various salt conditions at 298K. **(D)** $^1$H 1D NMR spectra of the imino region of 21-mer RNA indicating the imino and amino resonances (top panel) as identified by the $^1$H-$^{15}$N natural abundance SOFAST-HMQC (bottom panel) recorded on 500 µM sample in the presence of 5 mM KCl at 278 K. **(E)** Circular dichroism spectrum of 38.5 µM 21-mer RNA in 5 mM potassium phosphate buffer pH 6.5, 1mM β-mercaptoethanol acquired at 25°C. **(F)** Thermal denaturation (25–95 °C) of 38.5 µM 21-mer RNA in 5 mM potassium phosphate buffer pH 6.5, 1mM β-mercaptoethanol as monitored by CD signals at 265 nm at a 0.5 °C min$^{-1}$ temperature gradient.

#### Supplementary Figure 6

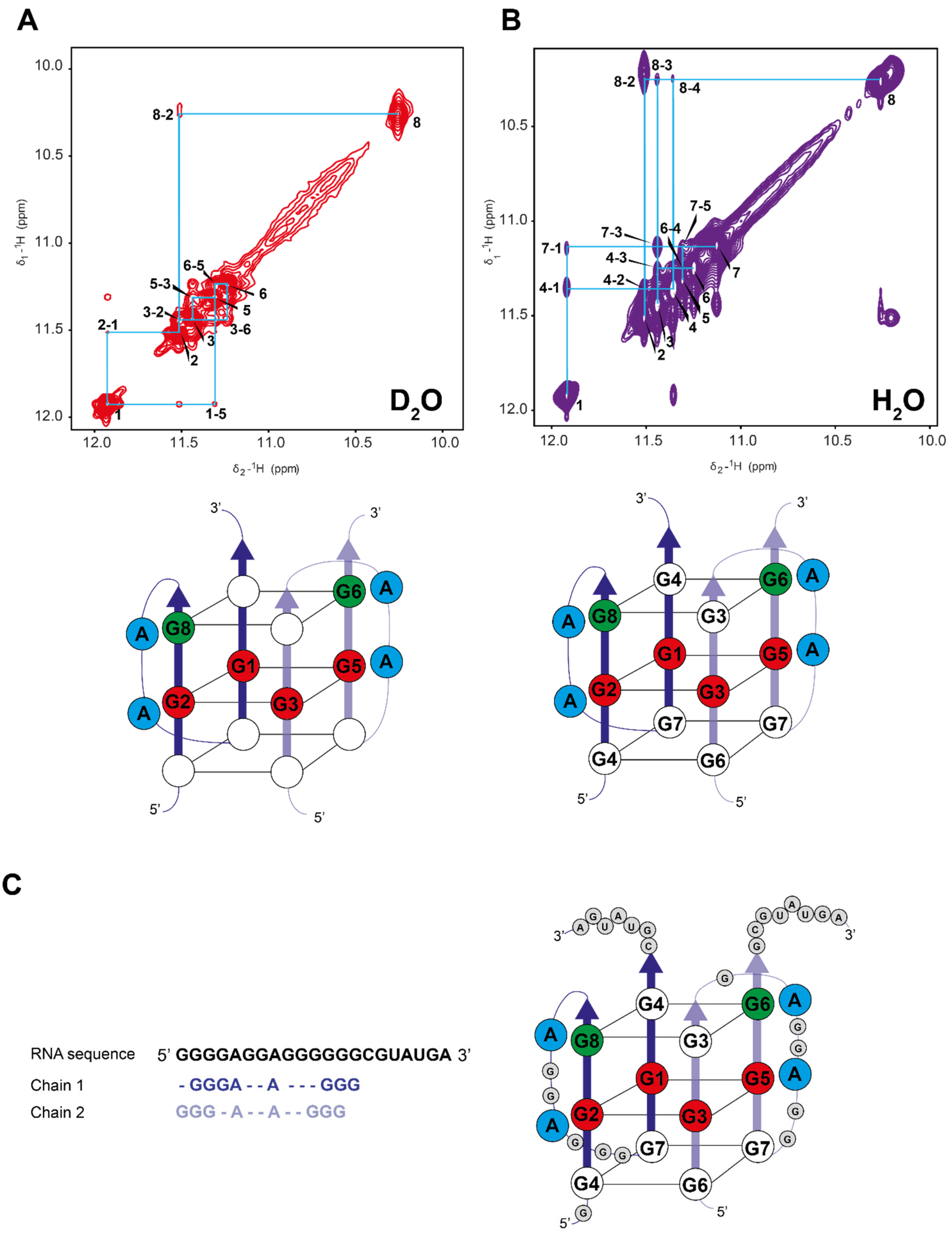

**Supplementary Figure 6. $^1H$-$^1H$ imino NOESY correlations showing three planar G4 fold of poly G RNA.** 2D imino proton NOESY of 500 µM 21-mer RNA in 5 mM KCl recorded at 278 K in $D_2O$ **(A)** and in $H_2O$ **(B)**, with cross-peaks of 21-mer RNA G4 resonances and tentative resonance assignments annotated on the G4 model color coded same as **Figure 2E**. **(C)** 21-mer RNA sequence represented with putative residues involved in the G4 structure formation (left) and the proposed G4 model (right). RNA Chain 1 and 2 are indicated in blue and light blue color, respectively.

#### Supplementary Figure 7

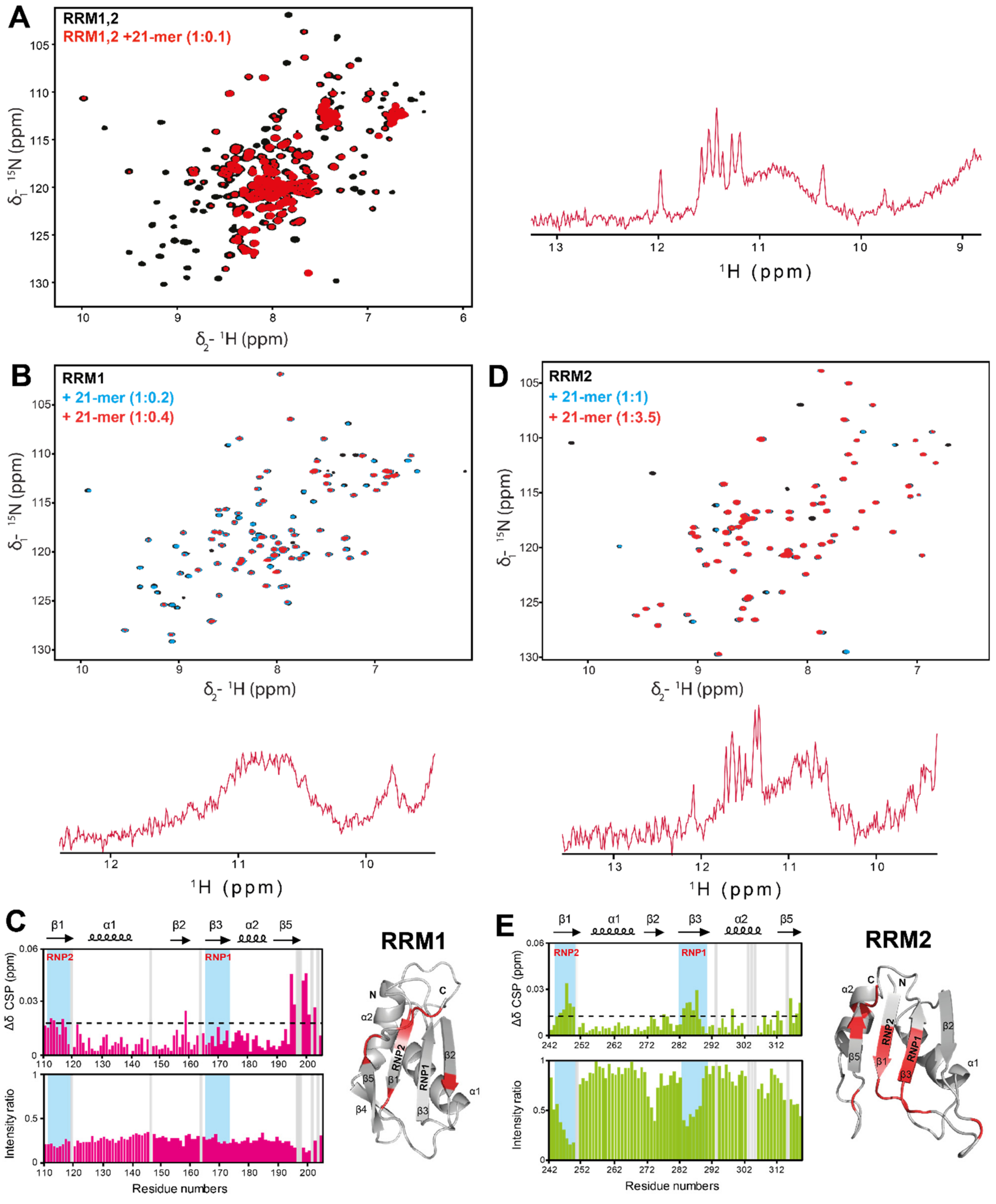


**Supplementary Figure 7. Characterization of LS2 RRMs and poly G RNA (21-mer) interaction. (A)** The overlay of the $^{1}H$-$^{15}N$ HSQC spectra of the LS2 RRM1,2 construct, in the absence (black) and in the presence of 0.1 molar excess (red) of the 21-mer RNA. **(B)** The overlay of the $^{1}H$-$^{15}N$ HSQC spectra of the LS2 RRM1 construct, in the absence (black) and in the presence of 0.2 molar excess (cyan) and 0.4 molar excess (red) of the 21-mer RNA. **(C)** Plots of chemical-shift perturbations (CSP, Δδ) and intensity change in the presence of 21-mer RNA with respect to residue numbers of RRM1. The secondary structure is shown at the top. Gaps in the plot caused by incomplete assignments or proline residues are indicated in grey bars. Both RNP sites are highlighted with a cyan background. Residues with CSPs higher than twice the standard deviation are mapped in red onto the structure of the RRM1. **(D)** The overlay of the $^{1}H$-$^{15}N$ HSQC spectra of the LS2 RRM2, in the absence (black) and in the presence of one molar excess (cyan) and 3.5 molar excess (red) of the 21-mer RNA. **(E)** CSP and intensity ratio plots showing changes in RRM2 chemical shifts and intensity in the presence of 21-mer RNA with respect to residue numbers of RRM2, respectively. The secondary structure is shown at the top. Both RNP sites are highlighted with a cyan background. RRM2 structure with residues undergoing CSPs higher than twice the standard deviation mapped in red.

#### Supplementary Figure 8

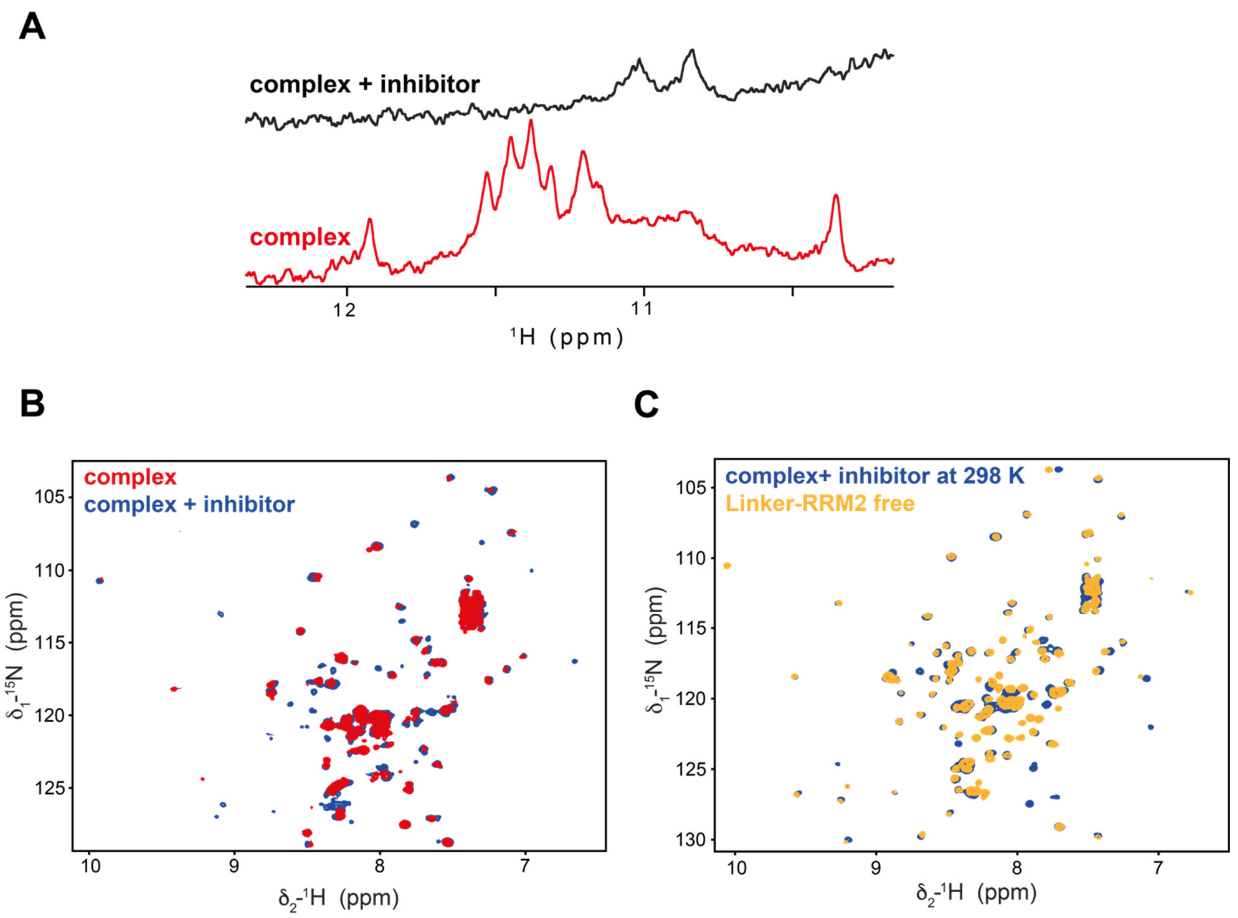


**Supplementary Figure 8. TmPyP4, a known G4 inhibitor abolishes the LinkerRRM2 and polyG RNA interaction. (A)** $^{1}$H 1D NMR spectra of the imino region of the linker-RRM2/21-mer complex in the absence (red) and in the presence of G4 specific inhibitor, TMPyP4 in nine molar excess (black) at 278 K. **(B)** The overlay of the $^{1}$H-$^{15}$N HSQC of the linker-RRM2-21-mer complex in the absence (red) and the presence of nine molar excess of the G4 specific inhibitor (black) at 278 K. **(C)** Comparison of the $^{1}$H-$^{15}$N HSQC spectrum of the linker-RRM2-21-mer complex in presence of nine molar excess after overnight incubation at RT (black) with $^{1}$H-$^{15}$N HSQC spectrum of free linker-RRM2 (cyan) at 298 K. Chemical shifts of the free protein match quite well (except residues of the linker helix) with the chemical shifts of the complex in the presence of inhibitor.

#### Supplementary Figure 9

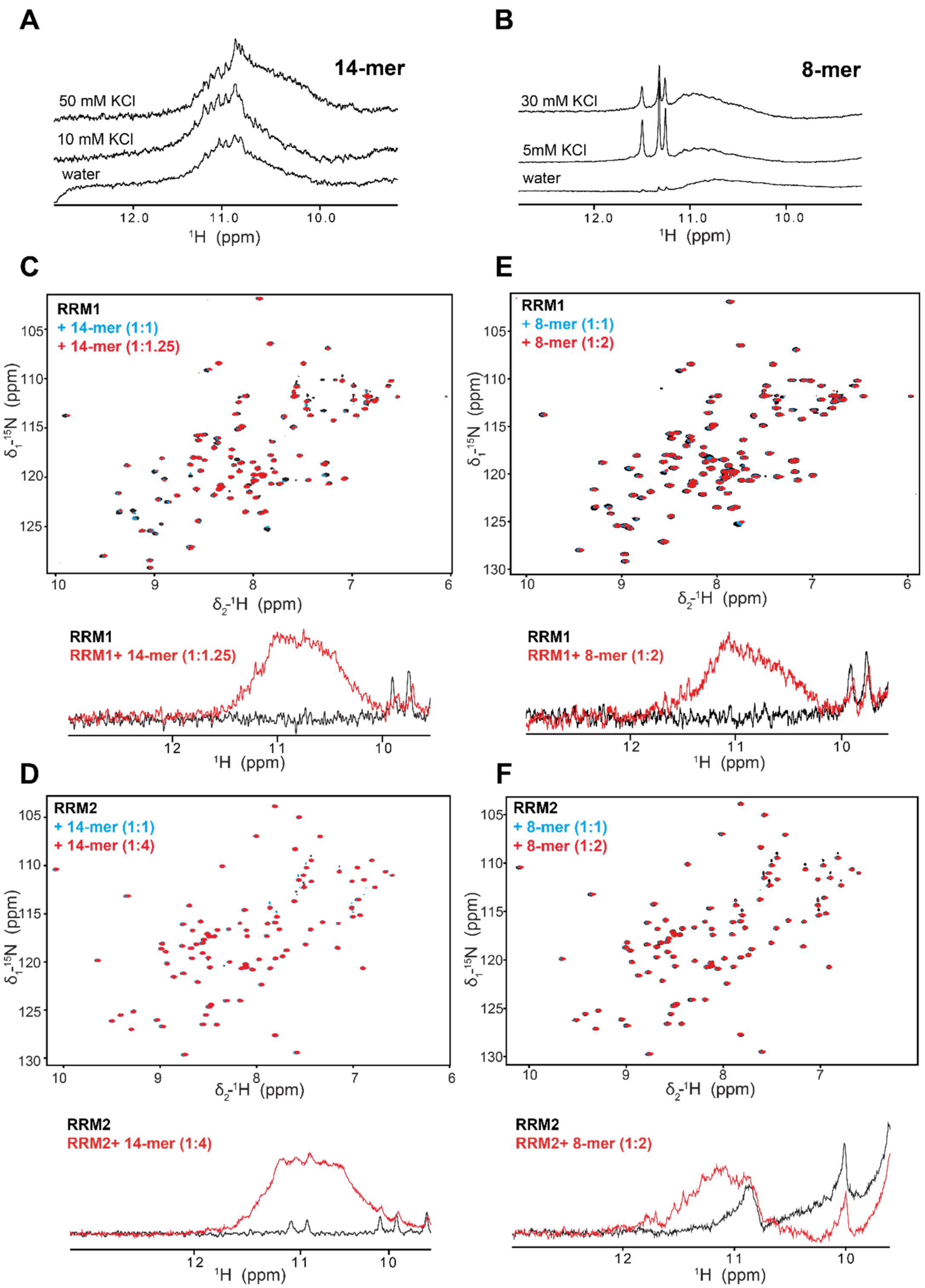


**Supplementary Figure 9. Characterization of Guanosine rich RNAs (14-mer and 8-mer) and their interaction with individual LS2 RRMs. (A)** $^{1}$H 1D NMR spectra of the imino region of the 14-mer RNA in different KCl concentrations shows that RNA exists mainly in heterogeneous population (broad peak around

11 ppm). **(B)** $^1$H 1D NMR spectra of the imino region of the 8-mer RNA adopts a uniform conformation in the presence of low KCl concentration and with the increase in the KCl concentration, heterogeneous population are also observed (broad peak around 11 ppm). The overlay of $^1$H-$^{15}$N HSQCs showing the comparison of the 14-mer RNA titration with RRM1 (**C**) and RRM2 (**D**) along with respective $^1$H 1D NMR spectra in the imino region showing G4 formation. The overlay of $^1$H-$^{15}$N HSQCs in the presence of 8-mer RNA with RRM1 (**E**) and RRM2 (**F**) respectively. $^1$H 1D NMR spectra of the imino region indicating the formation of G4 species are shown at the bottom.

#### Supplementary Figure 10

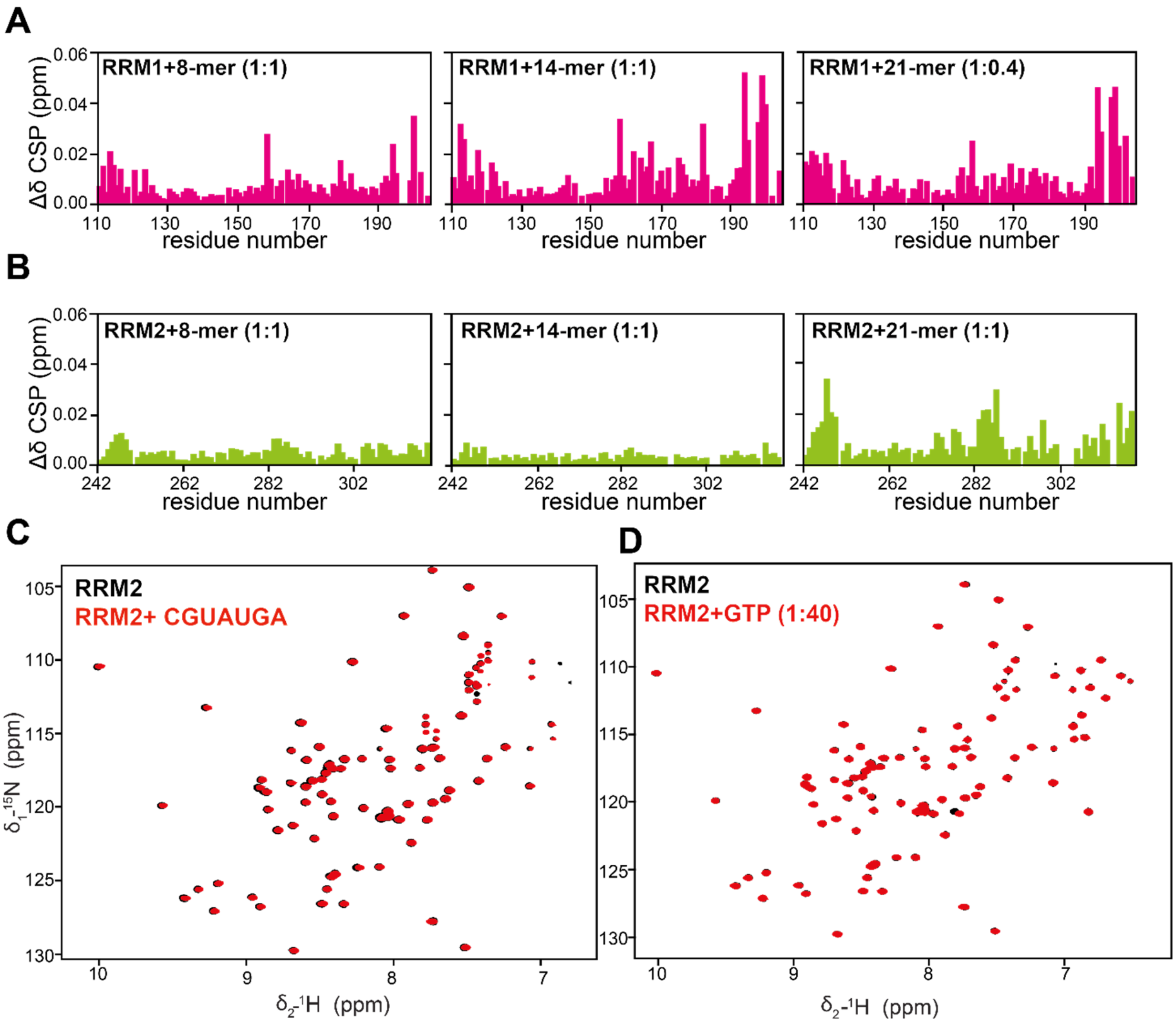


**Supplementary Figure 10. LS2 RRM2 shows specificity towards the poly-G 21-mer RNA. (A)** The comparison of CSP plots (Δδ) RRM1 and **(B)** RRM2 in presence of various guanosine-rich RNA oligonucleotides. RRM1 shows interaction with all three oligonucleotides whereas, RRM2 shows significant specificity towards 21-mer RNA. **(C)** The overlay of the $^{1}H$-$^{15}N$ HSQC of the LS2 RRM2 in the absence (black) and in the presence of CGUAUGA (red) and **(D)** GTP nucleotides (red) indicating lack of interaction.

#### Supplementary Figure 11

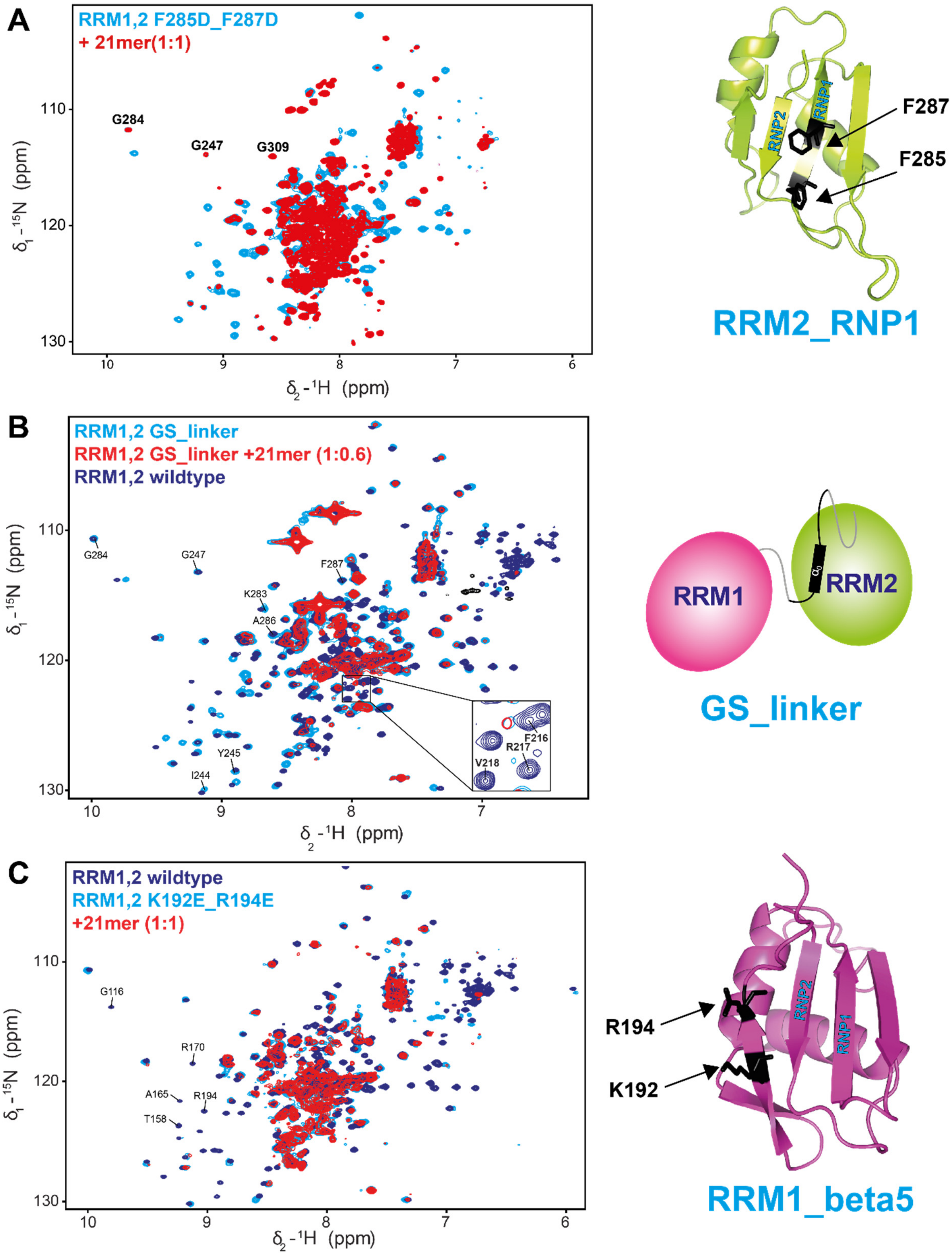

**Supplementary Figure 11. NMR analyses of the LS2 mutants designed to disrupt RNA binding contributions. (A)** The overlay of the $^{1}H$-$^{15}N$ HSQC of the RRM1,2 F285D_F287D double mutant in the absence (cyan) and in the presence of 21-mer RNA (red). Selected RNP residues of RRM2 involved in the RNA binding in the wild type protein are annotated. This double mutant protein has impaired RNA binding activity from the RRM2 domain, as seen by lack of effect of certain RRM2 residues, which show characteristic changes in the wild type protein in the presence of the RNA. Residues selected for the mutations (F285 and F287) are highlighted in black on the RRM2 structure. **(B)** The comparison of the $^{1}H$-$^{15}N$ HSQCs of the RRM1,2 GS linker mutant in the absence (cyan) and in the presence of the RNA (red) with the wild type protein (blue). Inset shows the missing $\alpha_0$ helix specific resonances from the RRM1,2 GS linker mutant protein. The comparison shows that the mutant protein is folded and also both RRM domains retain the RNA binding activities. **(C)** The overlay of the $^{1}H$-$^{15}N$ HSQC of the RRM1,2 K192E_R194E double mutant in the absence (cyan) and in the presence of 21-mer RNA (red) and wild type protein (blue). The mutation results in the unfolding of the RRM1 domain while retaining the RNA binding activity from linker and RRM2 domain. Few representative residues of RRM1 in the wild type protein are annotated for the comparison. The residues selected for the mutations (K192 and R194) are shown in black color on the RRM1 CS-ROSETTA structure.

#### Supplementary Figure 12

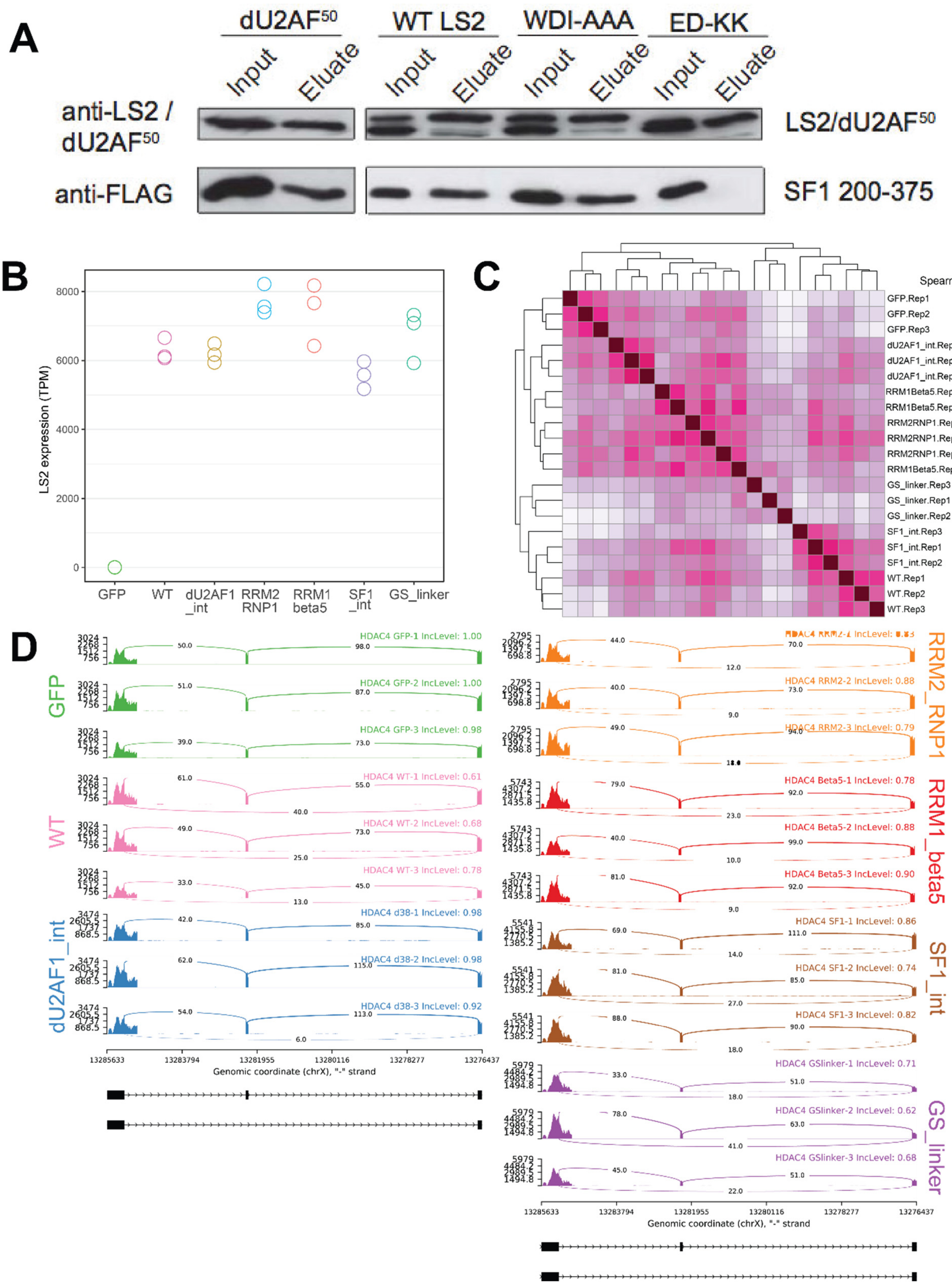

**Supplementary Figure 12. Biochemical characterization for LS2 mutants, RNA expression and alternative splicing analysis (A)** GST pulldown assay to assess interaction between LS2 and SF1. Purified recombinant GST tagged LS2 (or, as a control, dU2AF50) was incubated with a FLAG-tagged peptide corresponding to amino acids 200-375 of *Drosophila* SF1. Two different LS2 mutants were tested. WDI-AAA is a mutation of three residues required for interaction with dU2AF38 (W62, D63, and I64). ED-KK is a mutation of two residues (E370 and D371) that are predicted to be important for interaction with SF1 based on their homology to residues in hU2AF2 that are required for interaction with human SF1[5]. **(B)** RNA expression values (TPM) for LS2 in each mutant-expressing line. **(C)** PSI values for alternative exons that were significantly differentially included when comparing GFP samples to any LS2 sample were used to compare samples by Spearman correlation and hierarchical clustering. **(D)** Sashimi plot depicting the inclusion of an alternative exon that is strongly repressed by wildtype LS2 but displays less repression in the LS2 mutants.
